# Odorant receptor copy number change, co-expression, and positive selection establish peripheral coding differences between fly species

**DOI:** 10.1101/2021.09.19.460991

**Authors:** Thomas O. Auer, Raquel Álvarez-Ocaña, Steeve Cruchet, Richard Benton, J. Roman Arguello

**Affiliations:** Center for Integrative Genomics, Faculty of Biology and Medicine, University of Lausanne, Lausanne, Switzerland; Department of Ecology & Evolution, Faculty of Biology and Medicine, University of Lausanne, Lausanne, Switzerland; Swiss Institute of Bioinformatics

## Abstract

Despite numerous examples of chemoreceptor gene family expansions and contractions, how these changes relate to modifications in the neural circuitry in which they are expressed remains unclear. *Drosophila*’s Odorant receptor (Or) family is ideal for addressing this question because a large majority of Ors are uniquely expressed in single olfactory sensory neuron (OSN) types. Between-species changes in *Or* copy number, therefore, may indicate diversification/reduction of OSN populations. To test this, we investigated a rapidly duplicated/deleted subfamily (named *Or67a*) in *Drosophila melanogaster* and its sister species (*D. simulans, D. sechellia*, and *D. mauritiana*). We found that the common ancestor had three *Or67a* paralogs that had already diverged adaptively in their odor-evoked responses. Following their speciation, two *Or67a* paralogs were lost independently in *D. melanogaster* and *D. sechellia*, with ongoing positive selection acting on the intact genes. Instead of the expected singular expression of each diverged Ors, the three *D. simulans* Or67a paralogs are co-expressed. Thus, while neuroanatomy is conserved between these species, independent selection on co-expressed receptors has contributed to species-specific peripheral coding of olfactory information. This work reveals a model of adaptive change previously not considered for olfactory evolution and raises the possibility that similar processes may be operating among the largely uninvestigated cases of Or co-expression.

## 2. Introduction

The evolution of animal chemoreceptor families is characterized by rapid changes in gene copy number [1–6]. Numerous studies have correlated expansions and contractions of these families to known ecological shifts (i.e. dietary change, host-plant specialization), reflecting their capacity to quickly respond to environmental variation and to contribute to adaptive modifications [2, 7–10]. The evolution of chemoreceptor gene repertoires has raised considerable interest from a molecular evolution perspective, where they are often modeled as stochastic birth-and-death processes [11, 12]. These deletion and duplication events become additionally compelling in light of their roles in establishing the peripheral coding of chemical environments, and due to their extremely selective expression patterns: only one, or a small few, are expressed per neuron population. Thus, in addition to the gene family changes, understanding how chemoreceptor duplicates evolve cell-specific expression, and how chemoreceptor gains and losses functionally impact the sensory cells in which they are expressed, remain a puzzle.

A challenge to addressing these questions is the need for experimentally tractable systems with which to link changes at the level of the genome to physiology and neuroanatomy. *Drosophila melanogaster*, along with its closely related species, have emerged as an outstanding group for functional comparative studies of nervous systems. The extensive knowledge and resources that are available for *D. melanogaster* [13–16] is anchoring the development of genetic resources in its ecologically diverse sister species [17–21]. Additionally, the short evolutionary distances between multiple species in the *D. melanogaster* species group significantly aids in identifying key mutational events and in inferring the evolutionary processes that underly the changes of interest.

*Drosophila*’s Odorant receptor (Or) family is especially advantageous for relating between-species changes in chemoreceptor copy number to expression and neuronal response evolution. Odorant receptor expression is limited to the dendrites of neurons located in the fly’s two main olfactory organs, the antenna and maxillary palps. Importantly, other than a co-receptor (Orco [22]) expressed in all olfactory sensory neurons (OSN), a large majority of Ors are uniquely expressed in OSN populations (analogous to the singular expression of vertebrate odorant receptor genes [23, 24]). This distinct pattern of *Or* expression raises the intriguing possibility that between-species changes in *Or* copy number reflect the evolution of new OSN populations (in coordination with *Or* duplication events) or a simplification of the peripheral olfactory system (in coordination with *Or* deletion events). To investigate these questions, we focused on an *Or* subfamily named *Or67a*.

## 3. Results & Discussion

### 3.1 Or67a copy number is evolutionary dynamic

*Or67a* is one of the few olfactory receptor genes that differs in copy number among the closely-related *Drosophila melanogaster* species subgroup [2, 12, 25], which share a common ancestor *∼*3.4 million years ago [27]. It has also experienced remarkable expansions in more distantly-related species (for example, *D. suzukii*, which shares a last common ancestor with *D. melanogaster* ∼15 million years ago [28], has five paralogs (Fig. 1A)) [12, 25, 26]. Recently, *D. melanogaster* Or67a was shown to be the most broadly responding antennal Or when presented with headspace odors from an extensive collection of fruits [29], suggesting that evolution of the Or67a subfamily may be related to species-specific responses to food odors. To connect between-species differences in *Or* copy number and functional changes in the sensory neurons where they are expressed (Fig. 1B), we focused on *D. melanogaster* and its sister species in the *simulans* complex (*D. simulans, D. sechellia*, and *D. mauritiana*), which share a common ancestor *∼*0.24 million years ago [30, 31]. *D. sechellia* and *D. melanogaster* have a single intact *Or67a* gene while *D. simulans* and *D. mauritiana* have three (Fig. 1A). We refer to these three *Or67a* genes as *Or67a*.*P* (the 3L copy proximal to the centromere), *Or67a*.*D* (the 3L copy distal to the centromere), and *Or67a*.*3R* (the copy on the right arm of the third chromosome).

**Figure 1:**
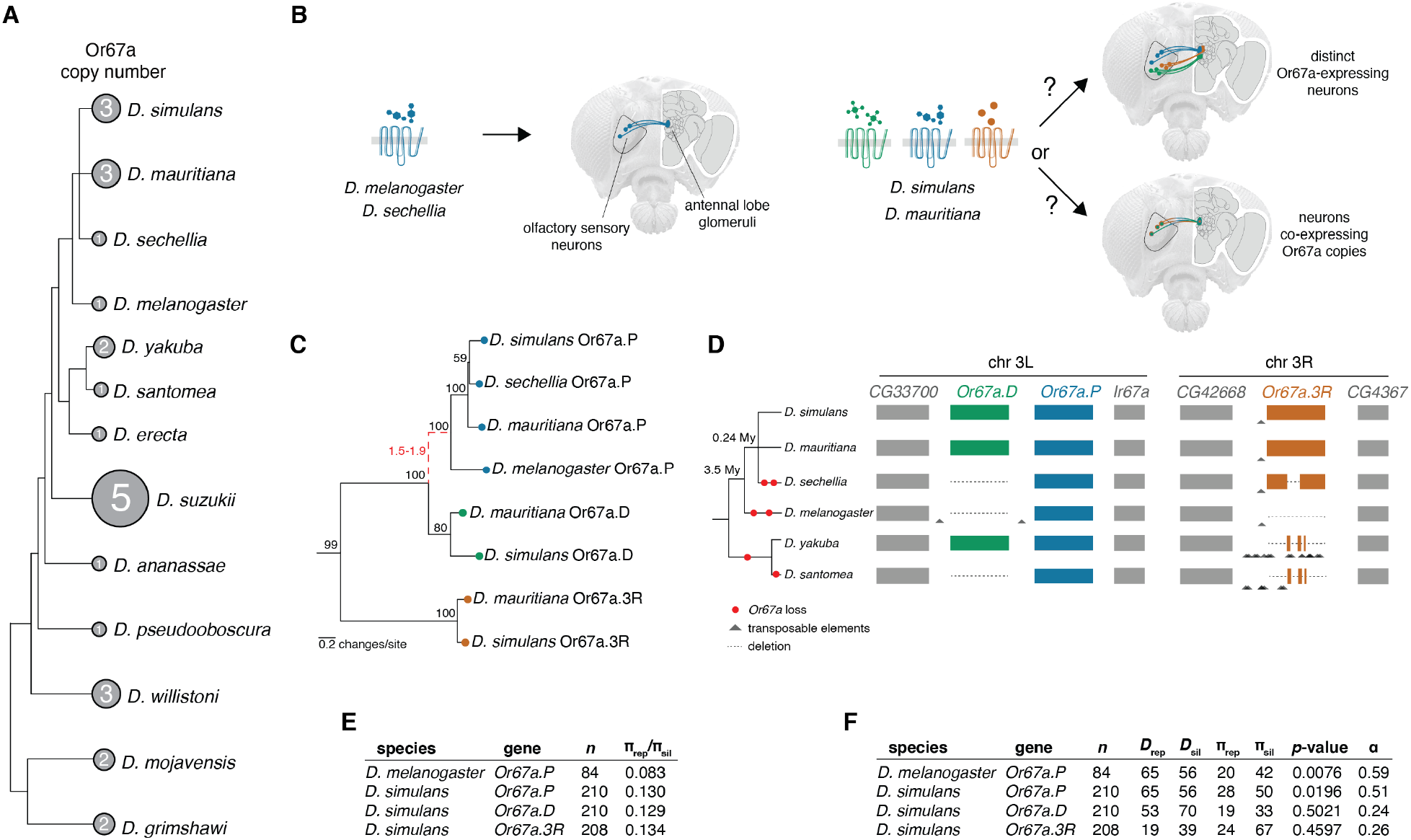
**(A)** *Drosophila* species tree (branches not to scale) illustrating *Or67a* copy number changes that have been reported in a subset of available genome assemblies (i.e. [2, 12, 25, 26]). These numbers exclude pseudogenes that are recognizable *Or67a* family members. **(B)** Illustration of the evolutionary scenario that is being investigated. To what extent are the three receptors tuned to different ligands? For *D. simulans* and *D. mauritiana*, are the three receptors expressed in distinct neuron populations that project to different regions (glomeruli) of the antennal lobe, or are they co-expressed in the same neuron population? **(C)** Bayesian protein tree inferred for the intact Or67a subfamily. Black numbers near nodes indicate posterior support. The branch with a dashed line was inferred to have a significant elevation in protein evolution (dN/dS = 1.5-1.9, red text); the remaining branches were inferred to have been under functional constraint (dN/dS < 0.5). **(D)** Overview of the parallel loss of the *Or67a.D* and *Or67a.3R* genes in *D. melanogaster* and the *simulans* group, using *D. yakuba* and *D. santomea* as outgroup species. On the left is a species tree for the samples included in the analyses (branches not to scale). Numbers at tree nodes indicate divergence dates in millions of years (My) for *D. melanogaster* and the *D. simulans* group [30, 31]. To the right of the tree are schematics of the alignments of the *Or67a*-containing chromosomal regions. Shown are the conserved genes (grey rectangles) that flank the *Or67a*-containing regions (colored rectangles) and the independent deletions of *Or67a.D* and *Or67a.3R* genes (dashed lines). The deletions are mapped onto the species tree with red dots. Many remnants of transposable elements were identified within these intervals, illustrated with grey triangles (the schematic is not to scale, but see Figs. S1, S2 and Files S1-3). **(E)** Table summarizing functional constraint on *Or67a* paralogs as measured by the ratio of nucleotide diversity at amino-acid replacement positions (π_*rep*_) to nucleotide diversity at silent positions (π_*sil*_). All copies have π_*rep*_/π_*sil*_ *<* 0.5, indicating ongoing purifying selection. Sample sizes are indicated by *n*. **(F)** Table summarizing McDonald-Kreitman tests for adaptive protein evolution and *α*, the estimated number of amino acid substitutions fixed by positive selection. The *Or67a.P* copies were found to have experienced adaptive protein evolution in both *D. melanogaster* and *D. simulans*, while signatures of adaptation were not found in *simOr67a.D* or *simOr67a.3R*. #*Drep* = number of amino acid replacement substitutions; #*D*_*sil*_ = number of silent substitutions; #π_*rep*_ = number of amino-acid replacement polymorphisms; #π_*sil*_ = number of silent polymorphisms. Sample sizes are indicated by *n*.

To elucidate the evolutionary history of this *Or67a* subfamily, we first inferred a protein tree for the eight receptors. The well-supported tree clusters each of the Or67a.P, Or67a.D, and Or67a.3R members together, indicating that the three paralogs existed prior to this group’s speciation events, and that the *Or67a*.*D* and *Or67a*.*3R* copies were lost independently along *D. melanogaster*’s and *D. sechellia*’s branches (Fig. 1C). The scenario involving multiple independent *Or67a*.*D* and *Or67a*.*3R* losses is further supported by inspecting alignments of the homologous chromosomal regions, and polarizing the changes using the outgroup species, *D. yakuba* and *D. santomea*. The genes flanking the *Or67a* paralogs are conserved across the six species, verifying that the chromosomal regions are homologous (Fig. 1D, Files S1,2). However, considerable nucleotide and indel differences have arisen between species within the intervals containing the *Or67a* paralogs, as have remnants of transposable elements, particularly for the *Or67a*.*3R*-containing region in *D. yakuba* and *D. santomea* (Fig. 1D and Fig. S1,2, File S3). These alignments clarify that independent deletions have completely removed the *Or67a*.*D* ortholog in *D. sechellia* (*secOr67a*.*D*), *D. melanogaster* (*melOr67a*.*D*), and *D. santomea*, although it remains intact in *D. yakuba*. Deletions have also entirely removed the *melOr67a*.*3R* ortholog, and a portion of the coding region in the *D. sechellia* ortholog (*secOr67a*.*3R*), eliminating sequences encoding two transmembrane domains that are required for forming the ion channel of the receptor [32, 33]. Short remnants of the *Or67a*.*3R* orthologs are still detectable in *D. yakuba* and *D. santomea*, additionally indicating that the ortholog was present in these more distant species. In combination, these data support a history in which three *Or67a* paralogs existed in the common ancestor of *D. melanogaster* and the *simulans* group, and that *D. sechellia* and *D. melanogaster* have recently lost the *Or67a*.*D* and *Or67a*.*3R* copies in parallel. The rapid change in *Or67a* copy numbers is likely related to past transposable element insertions and deletions in these loci.

### 3.2 Recurrent positive selection on Or67a paralogs

The observation of recent parallel gene losses among these closely-related species raises questions about the selective constraints acting on the intact receptors. We tested models of protein evolution by fitting rates of amino acid-changing (dN) and silent (dS) substitutions along the branches of the Or67a tree (Fig. 1C). Among the models we investigated, those that fit best consistently resulted in strong selective constraint along nearly all branches (dN/dS < 0.45). The branch leading to the Or67a.P clade was the only exception, with an elevated rate of amino acid changes that is consistent with positive selection acting on the Or67a.P coding sequence following the *Or67a.D*/*Or67a.P* tandem duplication event (dN/dS = 1.5-1.9; Tables S1 and S2, File S4). Using available population genomic datasets for *D. melanogaster* and *D. simulans*, we carried out more sensitive tests of ongoing purifying selection based on the ratio of amino acid to silent polymorphism (π_*rep*_/π_*sil*_). These measures lent additional support for functional constraint currently acting on the intact Or67a members for these two species, with all π_*rep*_/π_*sil*_ < 0.2 (Fig. 1E; File S5).

Combining our polymorphism datasets with between-species alignments, we applied McDonald-Kreitman tests of adaptive protein changes [34], and estimated the fraction of amino acid substitutions that have been fixed within a species by positive selection (*α*) [35]. These analyses also identified signals of adaptive protein evolution for the *Or67a.P* copies in both *D. melanogaster* and *D. simulans*, where 75% and 54%,respectively, of the protein changes were estimated to have been fixed by positive selection (Fig. 1F). In contrast, the *simOr67a.D* and *simOr67a.3R* copies did not carry signatures of adaptation. This *D. melanogaster* result is consistent with previous population genomic studies that identified *melOr67a.P* as evolving adaptively between species, as well as experiencing very recent positive selection between extant populations [36, 37], and further underscores past and ongoing adaptive changes in *melOr67a.P*. It has also been hypothesized that adaptive receptor gene loss may be an important route for sensory change within chemosensory systems [38]. If the two *D. melanogaster* deletions were adaptive and swept to fixation in the recent past, reduced genetic variation (and a negative Tajima’s *D* [39]) may be detectable [40]. However, analyses of the polymorphism at the loci containing these deletions did not provide evidence of adaptive loss, as the genetic variation was not different from the larger surrounding chromosomal intervals (Fig. S3).

These evolutionary genetic results provide evidence that the intact *Or67a* genes are currently under functional constraint, despite parallel gene losses in the recent history of the subfamily. They additionally highlight recurrent bouts of positive selection that presumably diversified receptor function, particularly for the *Or67a.P* clade.

### 3.3 Positive selection has diversified Or67a receptor tuning

To test the hypothesis that positive selection has contributed to the diversification of receptor function within the Or67a subfamily, we performed *in vivo* electrophysiological recordings of odor-evoked activity of Or67a paralogs and orthologs. We expressed individual Or67a receptors in a *D. melanogaster* antennal basiconic 3A (ab3A) “decoder” neuron, which lacks its endogenous receptor [41] (Fig. 2A,B), and quantified neuronal responses to a panel of nine odors. These nine odors were selected to cover a range of strong to weak Or67a.P ligands based on previous work in *D. melanogaster* [42–44]. Globally, we observed highly significant evolutionary changes in odor response profiles (global Wilks’ Lambda = 15.06, *p <<* 0.01; Fig. 2C). When testing for differences across the Or67a.P/D/3R paralogs within *D. mauritiana* or *D. simulans*, all comparisons were significantly different (*p* < 0.01). When testing for differences among orthologs across species (between all four species for Or67a.P or between *D. simulans* and *D. mauritiana* for Or67a.D and Or67a.3R), all comparisons were again significant (*p* < 0.01), except for the responses measured for Or67a.3R orthologs of *D. simulans* and *D. mauritiana* (*p* > 0.01). The statistical approach that we used to test for differences in odor responses [45] also allowed us to calculate the relative effects that the odors have on the receptor responses, thereby highlighting key odors combinations that drove these significant ortholog/paralog differences (Fig. 2D; Table S3). For example, the relative effect of geraniol on *sim*Or67a.3R is 86%, indicating a high probability of this receptor responding the strongest to this odor given a random sample from the full set of recordings (a comparable effect exists for *mau*Or67a.3R, 85%). Similarly, a recording from *sim*Or67a.D has a 97% probability of having the strongest response to ethyl hexanoate, which together with the large relative effects of pentanoic acid, methyl hexanoate, and *2*-heptanone, strongly differentiate it from its *mau*Or67a.D ortholog. Clustering these response data using principal component analysis (PCA) further highlights the evolutionary changes among receptors (Fig. 2E). In particular, the variation among Or67a.P copies is readily apparent, as is the distinct separation of the two Or67a.3R copies (together with *sec*Or67a.P) from the other receptors.

**Figure 2:**
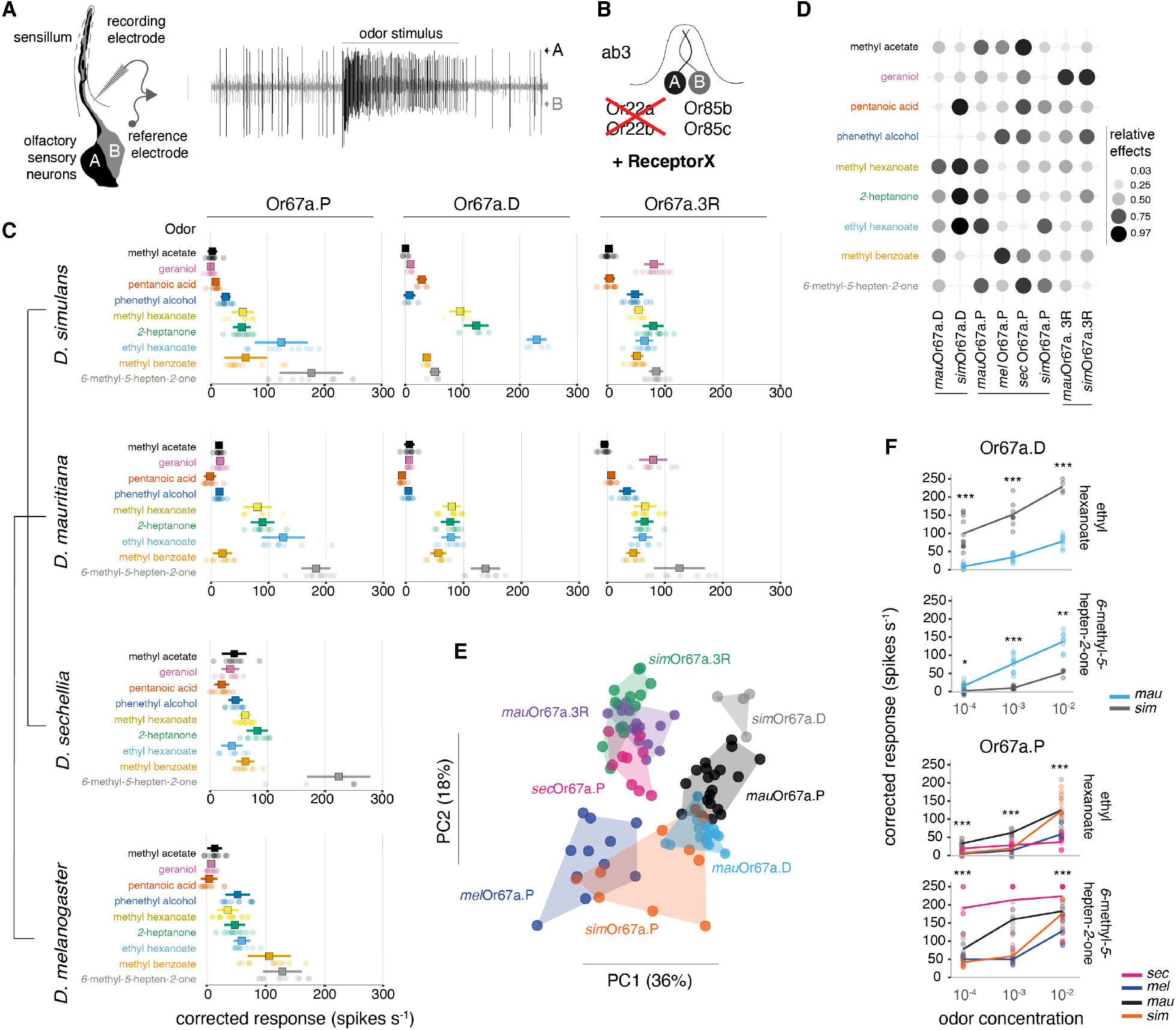
**(A)** Schematic for single sensilla recordings and a resulting spike train. The example displays a sensillum housing two olfactory sensory neurons (A and B), differentiable by large and small spike amplitudes. **(B)** Illustration of the *D. melanogaster* “decoder” system used to screen the *Or67a* from the four species [41]. **(C)** Quantification of Or67a.P/D/3R responses to a panel of nine odors at 10^−2^(v/v) concentration, organized by the species relationship (tree in left margin, not to scale). Squares indicate the mean and error bars display the standard deviation. Samples size (number of independent sensilla recorded from) per odor/receptor combination = 4 -12. **(D)** Relative effects of the odors on Or67a.P/D/3R responses at 10−2(v/v) concentration. These values provide the probability that a given odor-receptor response will be the largest given the full dataset. **(E)** Principal component analyses based on the data from Fig. 2B. Percentages along the axes indicate the amount of variation explained by the principal components. Species names have been abbreviated to the first three letters. **(F)** The two odor-receptor combinations that resulted in the largest dose-response differences among the Or67a.P/D/3R orthologs (see Fig. S4 for the other odors). For simplicity, the level of significance indicated above each concentration’s comparison is only for the single species comparison with the largest difference (see Table S4 for the full set of tests; \**p* < .05, \*\**p* < .01, \*\*\**p* < .001). *p*-value correction for multiple comparisons was done using the Holm method. Sample sizes = 4-11.

We tested for differences in sensitivity to our panel of odors by generating dose-response curves for the five odors that resulted in the strongest responses at the 10−2(v/v) concentration. These experiments revealed numerous differences in sensitivity among both paralogs and orthologs, but were most pronounced in Or67a.P and Or67a.D, concordant with the elevated diversity in response profiles to the full odor panel at the 10−2 concentration (Figs. 2F, S4). For example, *sim*Or67a.D is significantly more sensitive across concentrations of ethyl hexanoate than *mau*Or67a.D (*p* < 0.01; Table S4), while the opposite is the case for *6*-methyl-*5*-hepten-*2*-one (*p* < 0.01; Table S4). Other notable differences are the Or67a.P responses to *6*-methyl-*5*-hepten-*2*-one, where *sec*Or67a.P has high sensitivity across all concentrations, with additional species differences increasing with concentration (Fig. 2F; Table S4). These data demonstrate widespread evolution of ligand response profiles within the Or67a subfamily, supporting our molecular evolutionary inferences that positive selection has contributed to functional changes.

### 3.4 The three D. simulans Or67a paralogs are co-expressed

Our evolutionary genetic analyses and electrophysiological experiments uncovered adaptive functional diversification in the Or67a subfamily. The three *D. simulans* receptors could either define different populations of OSNs (two of which were lost in *D. melanogaster* and *D. sechellia*) or be co-expressed in a single neuron population (Fig. 1B). To investigate these possibilities, we first examined receptor expression using RNA fluorescence *in situ* hybridization (FISH). For all of the four species, we detected Or67a-expressing neurons within a comparable spatial domain of the antenna (Fig. 3A). Quantification of neuron numbers indicates a similar number of cells expressing *Or67a.P* and *Or67a.D* in *D. simulans*. However, the high sequence identity of these genes (Table S5) might result in cross-hybridization of RNA probes. We observed very few *Or67a.3R* positive cells (possibly because of lower expression levels of this receptors); similarly, *mauOr67a.P* and *mauOr67a.3R* expression was weak but detectable.

**Figure 3:**
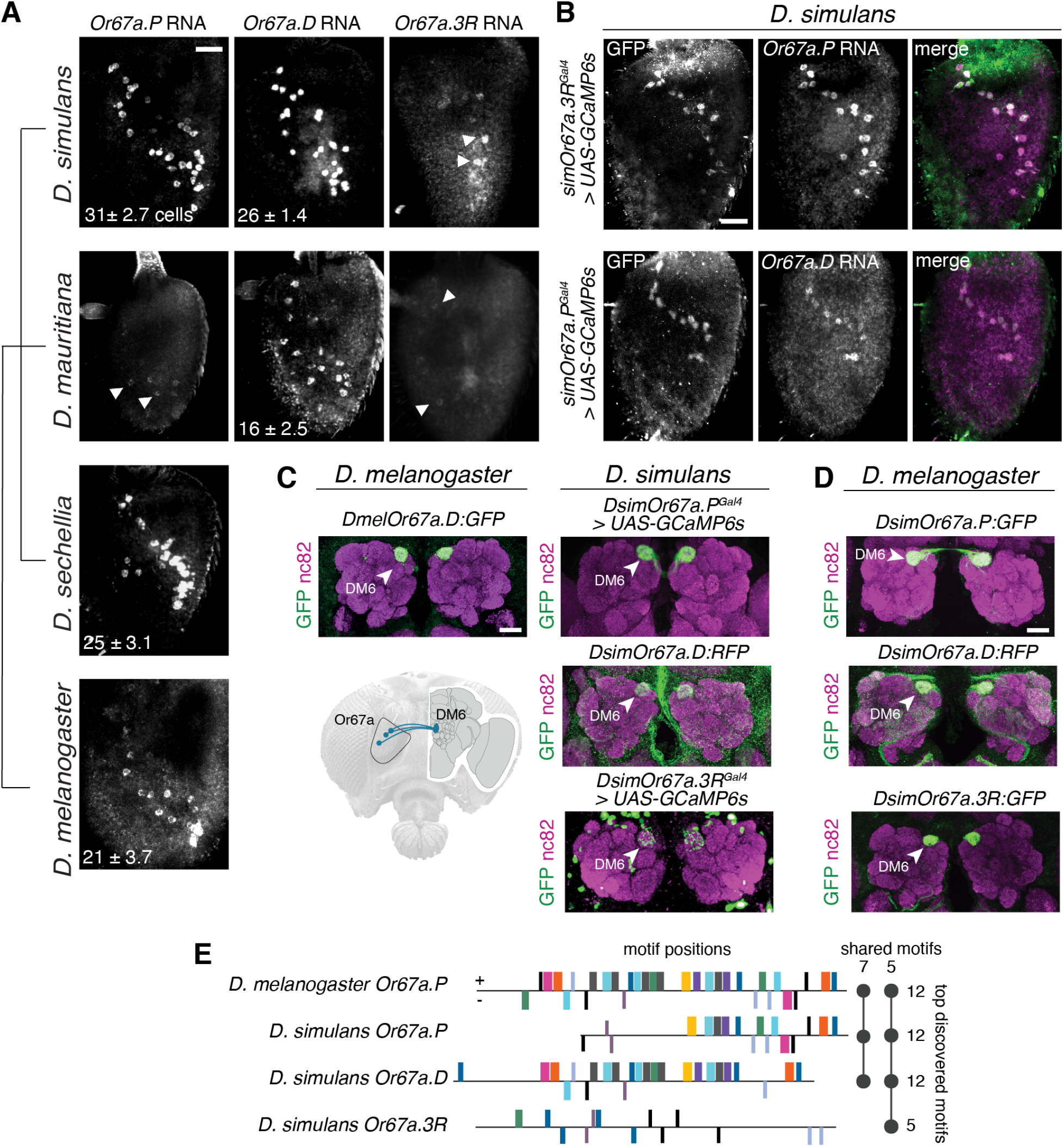
**(A)** Whole-mount antennal RNA expression of *Or67a* paralogs and orthologs in *D. simulans, D. mauritiana, D. sechellia* and *D. melanogaster* (top to bottom). Scale bar = 25 *µ*m. The number of *Or67a* expressing OSNs (+/- standard deviation) is indicated at the bottom left corner (n=5-12 antennae). Weak staining prevented quantification of OSN numbers expressing *Or67a.3R* in *D. simulans* and *Or67a.3R* and *Or67a.D* in *D. mauritiana*. Arrows point towards weakly labeled cells. **(B)** Antennal co-expression of knock-in Gal4 transcriptional reporters (visualized by *UAS-GCaMP6s*) and *Or67a.P* (top) and *Or67a.D* (bottom) RNA in *D. simulans*. Scale bar = 25 µm. **(C)** (left) Antennal lobe innervation of the promoter transcriptional reporter for *Or67a* in *D. melanogaster*. (below) Schematic illustrating the innervation of DM6 by Or67a-expressing neurons. (right) Gal4- and promoter transcriptional reporters for Or67a paralogs in *D. simulans*. All reporters label neurons innervating the DM6 glomerulus (arrowheads). Scale bar = 25 µm. **(D)** Antennal lobe innervation of transcriptional reporters for all three *D. simulans* paralogs in *D. melanogaster* (arrowheads show DM6 glomerulus). Scale bar = 25 *µ*m. **(E)** Putative regulatory motifs identified in the 5’ DNA sequences of the *Or67a* paralogs in *D. simulans* and *melOr67a.P* (1.5-2 kb, see methods). Boxes indicate the placement of candidate motifs, with colors illustrating the same motif sequence. Positive strand motifs are above the horizontal line and negative strand motifs are below. The sequences have been arranged to approximate a DNA alignment without gaps. The plot to the right summarizes the number of motifs per sequence and the overlap of motifs between the four sequences.

The high sequence similarity across paralogs, and the potential cross-reactivity of probes, prevents an un-ambiguous interpretation of paralog-specific cell number, as well as the use of double FISH for co-expression experiments. Therefore, we generated paralog-specific transgenic transcriptional reporters in *D. simulans*. We used a CRISPR/Cas9 mediated strategy to integrate Gal4 at the *simOr67a.3R* and *simOr67a.3P* loci (Fig. S5A), and combined both with a fluorescent reporter (UAS-GCaMP6s) to visualize promoter activity. Our attempts to generate an equivalent *simOr67a.D* Gal4 insertion were unsuccessful, so we generated a transgenic reporter for this gene using the upstream sequence of *simOr67a.D* to drive RFP expression (similar to a previous *melOr67a.P* reporter [46], Fig. S5B). Using these tools, together with RNA FISH, we confirmed that transcription from the *simOr67a.3R* locus overlaps with *simOr67a.P*, and transcription from the *simOr67a.P* locus overlaps with *simOr67a.D* mRNA expression (Fig. 3B). In the antennal lobe, *melOr67a.P* axonal projections uniquely innervate the DM6 glomerulus [46] (Fig. 3C). Similarly, both *D. simulans* Gal4 alleles (*simOr67a.P*^*Gal*4^ and *simOr67a.3R*^*Gal*4^), as well as the *simOr67a.D* transgenic reporter, uniquely labeled neurons targeting DM6 in *D. simulans* (Fig. 3C). These results collectively argue for the co-expression of the three *D. simulans* Or67a receptor paralogs in the homologous neuron population to *D. melanogaster Or67a* neurons.

The evolutionary stability of the co-expression of *D. simulans* paralogs is notable, given the divergence between their putative regulatory regions (Table S6). To investigate the transcriptional activity of these sequences outside of their endogenous genomic context, we generated transgenic transcriptional reporters containing the upstream sequences of each *D. simulans Or67a* paralogs and placed them in a common *D. melanogaster* background. Within the antenna, all three reporters display expression patterns that were consistent with the endogenous *melOr67a.P* mRNA (Fig. S5B, C), and they also paired within ab10 sensillum with Or49a/Or85f-expressing neurons (Fig. S5D). Moreover, all three labeled neurons target DM6 (Fig. 3D). Computational searches for putative regulatory motifs identified diverse patterns of overlap within the upstream sequence of the *Or67a* genes in *D. melanogaster* and *D. simulans*. Consistent with their sequence identity (Table S6), more motifs were shared between the upstream sequences of *Or67a.P* and *Or67a.D* than either shared with *Or67a.3R* (Fig 3E; Table S7). This observation suggests that co-expression of the three receptors has been maintained by diverse, possibly paralog-specific, regulators of Or expression.

### 3.5 D. simulans Or67a paralogs provide unique and overlapping contributions to peripheral tuning

The observation that the three *Or67a* paralogs are co-expressed in *D. simulans* led us to investigate the correspondence between the decoder neuron responses (Fig. 2B,C) and those from endogenous neurons (housed in ab10 sensilla) in *D. melanogaster* and *D. simulans*. For *D. melanogaster*, the wild-type ab10 response profile to the nine odors was qualitatively similar to that obtained from the decoder neuron experiments (Figs. 4A, 2C). In *D. simulans*, we observed the combined responses seen from three receptors that were individually expressed in the decoder neuron (though with overall lower responses). For example, the geraniol response that was seen only for the *sim*Or67a.3R paralog in the decoder neuron recordings was observed in the *D. simulans* wild-type ab10 responses. Additionally, ethyl hexanoate, which evoked the strongest responses across the three *sim*Or67a paralogs in the decoder neuron recordings, remained the strongest ligand in the *D. simulans* wild-type ab10 neuron (Fig. 4A). To investigate individual contributions of the *D. simulans* Or67a paralogs to the overall response profile in their endogenous neuron, we employed our *simOr67a.3R*^*RFP*^ and *simOr67a.P*^*Gal4*^ loss-of-function alleles. The recordings from the *simOr67a.3R*^*RFP*^ line revealed the loss of its unique response to geraniol and a significant reduction in its response to phenethyl alcohol (both Wilcoxon rank sum tests *p* < 0.01), but no modification in the responses to the other odors, consistent with *sim*OR67a.3R contributing uniquely to the global response profile (Fig. 4B; one-way MANOVA *F* = 5.45, *p* > 0.05; only the pairwise Wilcoxon rank sum tests for geraniol and phenethyl alcohol were significant). Based on the decoder neuron recordings, *simOr67a.P* had the highest responses to *6*-methyl-*5*-hepten-*2*-one compared to the other two paralogs. However, recordings from the *simOr67a.P*^*Gal4*^ flies did not result in a reduction in *6*-methyl-*5*-hepten-*2*-one response (Wilcoxon rank sum test p > 0.05), likely because *sim*Or67a.D and *sim*Or67a.3R both respond to this odor. Recordings from this mutant did not impact the global responses to the full panel of odors either (Fig. 4B; one-way MANOVA *F* = 2.42, *p* > 0.05), indicating functional overlap between *sim*Or67a.D and the other two Or67a paralogs (for at least these nine odors). Together, the results of these electrophysiological experiments highlight both specific and overlapping contributions that the Or67a paralogs make to the overall response profile of their endogenous OSNs.

**Figure 4:**
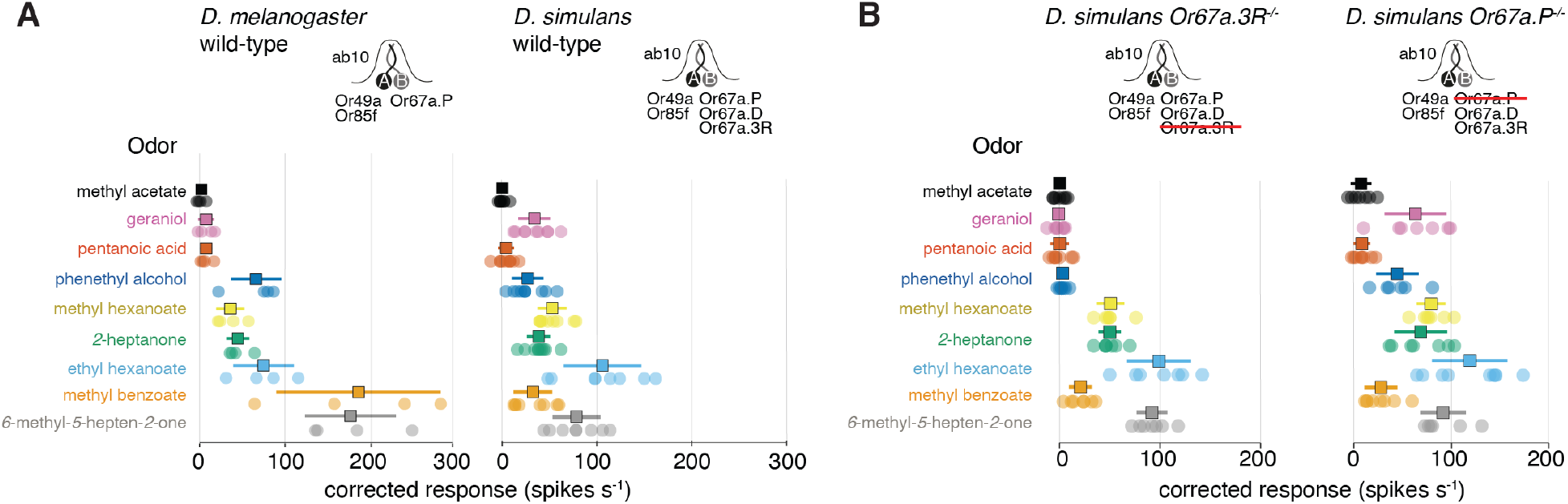
**(A)** Quantification of wild-type ab10 sensilla recordings for *D. melanogaster* (left) and *D. simulans* (right) to a panel of nine odors (as in Fig. 2C). Sample size = 4-10. **(B)** (right) Quantification of *simOr67a.3R* KO responses to the panel of nine odors. Sample size = 6. (left) Quantification of *simOr67a.P* knockout responses to the panel of nine odors. Samples size = 6-9. For both panels, squares indicate the mean, error bars display the standard deviation, and sample sizes refer to the number of independent sensilla recorded per odor/receptor combination.

## 4. Conclusions

Comparative electrophysiological analysis of odor responses of homologous neurons across species have identified many instances of evolutionary change [18, 19, 47–50]. Such changes are generally assumed to be due to modifications in the tuning of singularly expressed receptors, which have been supported by direct examination of the receptor responses in heterologous expression systems, and, in a few cases, the mapping of amino acid substitutions that underlie the differences between orthologous receptors [18, 19]. Our evolutionary study of the *Or67a* subfamily reveals an alternative mechanism in which positive selection can diversify olfactory receptors that remain co-expressed over millions of years, thereby providing additional degrees of freedom for a single OSN population to evolve novel peripheral tuning. This evolutionary mechanism is unlikely to be specific to the *Or67a* subfamily as copy number variation for other olfactory receptor subfamilies exists, as do several cases of odorant receptor co-expression (beyond Orco) [51–54]. For example, another fruit odor receptor, Or22a, and its paralog, Or22b, are co-expressed in *D. melanogaster* and have been shown to vary in copy number between *Drosophila* species [2, 12, 18, 41, 55, 56]. Additionally, the highly divergent Or33c and Or85e receptors are co-expressed in several fly species [54]. While physiological data suggest that some of these examples of co-expression impact neuron response properties [18, 47, 56, 57], more detailed evolutionary and expression studies - as presented for the Or67a subfamily - are needed to determine if similar processes are shaping other olfactory channels. The co-expression of multiple differentially-tuned receptors in a single neuron population is reminiscent of a widespread coding principle in the insect gustatory system, where it is common for combinations of co-expressed taste receptors to determine the tuning profile of gustatory sensory neurons [58–66]. If additional examples of olfactory receptor co-expression are shown to be evolutionarily stable, co-expression may be a feature more broadly shared between the olfactory and gustatory systems than previously appreciated.

## 5. Methods

### 5.1. Drosophila stocks

*Drosophila* stocks were maintained on standard wheat flour/yeast/fruit juice medium under a 12h light:12h dark cycle at 25°C. For *D. sechellia* strains, a few g of Formula 4-24^®^ Instant *Drosophila* Medium, Blue (Carolina Biological Supply Company) soaked in noni juice (nu3 GmbH) were added on top of the standard food.

### 5.2. CRISPR/Cas9-mediated genome engineering

#### sgRNA expression vectors

To express multiple sgRNAs from the same vector backbone, oligonucleotide pairs (Table S8) were used for PCR and inserted into *pCFD5* (Addgene no. 73914) via Gibson Assembly, as described [67]. For single sgRNA expression, oligonucleotide pairs (Table S8) were annealed and cloned into BbsI-digested *pCFD3-dU6-3gRNA* (Addgene no. 49410), as previously described [68].

#### Donor vectors for homologous recombination

Homology arms (1-1.6 kb) for *simOr67a.3R* were amplified from *D. simulans* (DSSC 14021-0251.195) genomic DNA and inserted into *pHD-DsRed-attP* [69] via restriction cloning. Oligonucleotide sequences are listed in Table S8. For endogenous tagging of *D. simulansOr67a.P* we generated a *T2A-Gal4* targeting vector flanked by homology arms (1-1.1 kb) via gene synthesis (GenScript Biotech) as described [70].

### 5.3. Molecular cloning and sequencing

#### UAS-cDNA vectors

To express the different Or67a receptors in the decoder neuron system, open reading frames were amplified from genomic DNA of the respective species via PCR, digested with restriction enzymes (BglII, EcoRI and/or KpnI) and integrated into *pUAST-attB* [71]. Oligonucleotide sequences are listed in Table S8.

#### OrX-reporter vectors

Promoter fragment for transcriptional reporters were amplified from *Dsim* (DSSC 14021-0251.195) genomic DNA via PCR, inserted into *pDONR221-MCS* [18] via restriction cloning and the resulting vector was combined with *pDEST-HemmarG* or *pDEST-HemmarR* [72] via LR recombination (Gateway, Thermo Fisher Scientific). Oligonucleotide sequences are listed in Table S8.

The oligonucleotides used for Sanger sequencing of *D. simulans* paralogs from multiple strains are listed in Table S8. The fasta sequences for these samples are found in Files S9-11.

### 5.4. Drosophila microinjections

Transgenesis of *D. simulans* and *D. melanogaster* was performed in-house following standard protocols, except for *simOr67a.D-RFP* transgenics (generated by Rainbow Transgenic Flies Inc). For CRISPR/Cas9-mediated homologous recombination, we injected a mix of an sgRNA-encoding construct (150 ng *µ*l-1), donor vector (400 ng *µ*l-1) and *pHsp70-Cas9* (400 ng *µ*l-1) (Addgene #45945) [69]. Site-directed integration into attP sites was achieved by co-injection of an attB-containing vector (400 ng *µ*l-1) and *pBS130* (encoding phiC31 integrase under control of a heat shock promoter (Addgene #26290) [73]). All concentrations are given as final values in the injection mix.

### 5.5. Electrophysiology

Single sensillum electrophysiological recordings were performed as described previously [74] using chemicals of the highest purity available from Sigma-Aldrich. Spike visualization and quantification was performed using AutoSpike32 (Syntech). To target ab10 sensilla in *D. melanogaster*, we used (*R*)-actinidine, which is a diagnostic odor for the neighboring Or85f-expressing neuron [75]. To target ab10 sensilla in *D. simulans*, we used fluorescent-guided recordings [76]. Spike visualization and quantification for these data performed using the Spike2 software (CED). Generally, we observed a lower response rate in *D. simulans* ab10 recordings compared to the recordings from the individually expressed receptors in the *D. melanogaster* “decoder” ab3A decoder neurons (Fig. 4B). This might be related to differences between the two recording rigs used for the experiments, but may also reflect a biological difference between natively-expressed and the misexpressed receptors. Odorants (*6*-methyl-*5*-hepten-*2*-one (CAS 110-93-0), methyl benzoate (CAS 93-58-3), ethyl hexanoate (CAS 123-66-0), *2*-heptanone (CAS 110-43-0), methyl hexanoate (CAS 106-70-7), phenethyl alcohol (CAS 60-12-8), pentanoic acid (CAS 109-52-4), geraniol (CAS 106-24-1), methyl acetate (CAS 79-20-9)) were used at 10−2(v/v) in all experiments (unless noted otherwise in the figures or figure legends) and diluted in paraffin oil or double distilled water. Corrected responses were calculated as the number of spikes in a 0.5 s window at stimulus delivery (200 ms after stimulus onset to take account of the delay due to the air path) subtracting the number of spontaneous spikes in a 0.5 s window 2 s before stimulation, multiplied by two to obtain spikes *s*^−1^. The amplitude of the A and B spikes in *D. simulans*’ ab10 did not differ greatly, and when the A cell fired upon odor stimulus the amplitude would “pinch” such that spike sorting by amplitude was not possible. As a result, the number of spikes for these recordings included both cells during the 0.5 s stimulation window. Odors that resulted in saturated bursts of spiking that were too numerous to count were replaced with the maximum value from those that were countable. The solvent-corrected responses shown in the figures were calculated by subtracting from the response to each diluted odor the response obtained when stimulating with the corresponding solvent. Recordings were performed on a maximum of three sensilla per fly. Response data was plotted using within R’s (v4.1.0 [77]) ggplot2 library (v3.3.0 [78]). To test for differences between Or67a.P/D/3R responses, we carried out a nonparametric multivariate approached implemented in the npmv library (v2.4, [45]) in R (see GitLab page). Principal component analyses were carried out with the “prcomp” function with in the R’s (v4.1.0) “stats” library, and plotted with the “scatterplot3d” library (v0.3.41 [79]). Missing data was imputed using the nonparametric approach implemented in R’s missForest (v1.4 [80]) on a per-odor basis. The full odor response datasets for all SSR experiments are provided in Files S6-8, and an R markdown file with analyses and plotting code are provided on our GitLab page.

### 5.6. Immunohistochemistry

RNA fluorescent *in situ* hybridization using digoxigenin- or fluorescein-labelled probes and immunofluorescence on whole-mount antennae were performed essentially as described [81, 82] using a rabbit *α*-GFP 1:500 (Invitrogen) and a chicken *α*-GFP 1:500 (Abcam) polyclonal antibody. *D. simulans* probe templates were generated by amplification of regions of genomic DNA (DSSC 14021-0251.004) using primer pairs listed in Table S8; these were cloned into *pCR-Blunt II-TOPO* and sequenced. Species specific *in situ* probes were generated for *D. melanogaster, D. sechellia* and *D. mauritiana* but did not show improved staining quality compared to *D. simulans* probes (data not shown). Immunofluorescence on adult brains was performed as described [83] using mouse monoclonal antibody nc82 1:10 (Developmental Studies Hybridoma Bank), rabbit *α*-GFP 1:500 (Invitrogen) and chicken *α*-GFP 1:500 (Abcam). Alexa488- and Cy5-conjugated goat *α*-rabbit and goat *α*-mouse IgG (Molecular Probes; Jackson Immunoresearch) and Alexa488-conjugated goat *α*-chicken (Abcam) secondary antibodies were used at 1:500.

### 5.7. Image acquisition and processing

Confocal images of antennae and brains were acquired on an inverted confocal microscope (Zeiss LSM 710) equipped with an oil immersion 40X objective (Plan Neofluar 40X Oil immersion DIC objective; 1.3 NA), unless stated otherwise. Images were processed in Fiji [84]. OSN numbers were counted using the Cell Counter Plugin in Fiji or Imaris (Bitplane).

### 5.8. Molecular evolution and polymorphism analyses

To infer the protein tree, Or67a.P/D/3R amino acid sequences were aligned using Clustal Omega with default settings [85]. The Or67a protein tree was inferred using Mr.Bayes (v3.2.7a) with the following settings: lset nucmodel=protein, mcmc nchains=6 ngen=10000, samplefreq=500, printfreq=100, diagnfreq=1000, burnin=500) [86]. To estimate dN/dS ratios over the branches of the Or67a subfamily tree, we used Maximum likelihood estimation implemented in PAML’s CODEML (v4.8 [87]), using the pamlX GUI (v1.3.1 [88]). Model testing was carried out using likelihood ratio tests on the outputted likelihoods of the models found in Table S1. For analyses of *D. simulans* polymorphism data in Fig. 1, we used an existing dataset [89], and the sequences from 15 additional strains (above). For the data set of Signor et al., we extract *Or67a.P/D/3R* regions from the full VCF file using VCFtools (v0.1.17 [90]), requiring a minimum mean depth of 5 (–min-meanDP 5) and sites that have a proportion of missing data greater that 0.5 (–max-missing 0.5). We converted these gene region VCF files to fasta format using the custom “vcf2fasta_remove_het.py” script, where nucleotides at heterozygous positions were sampled randomly. These fasta sequences were combined with the 15 sanger sequenced samples for the results shown in Fig. 1E,F. For *melOr67a.P*, we extracted the gene region for the prefilter VCF provided in [91]. For calculating silent and replacement diversity estimates, we used a custom script “calc_N_S.py” together with the paralog specific GTF file. Similarly, for silent and replacement divergence, we used a custom script “Div_N_S.py”. The custom scripts can be found our GitLab page. The reference genome used to make the alignments in Fig. 1D were: *D. melanogaster* v6.4 from flybase.org, *D. sechellia* and *D. simulans* from [92], *D. mauritiana* and *D. yakuba* from [93], and *D. santomea* from Prin_Dsan_1.0. The alignments of the *Or67a*-containing regions was generated with Clustal Omega (v1.2.3 [85]). Annotations of the transposable element fragments used RepeatMasker (v4.1.2-p1 [94]), with Dfam_3.0 and rmblastn v(2.9.0+), and existing annotations within flybase’s JBrowse.

### 5.9. Regulatory motif searches

We used the “MEME” programs within the MEME package (v5.4.1) to search for putative regulatory motifs within 5’ promoter regions of the *D. simulans* and *D. melanogaster Or67a* copies [95, 96]. We inputted 2kb for each gene, except for *simOr67a.P*, where only *∼*1.5 kb exists between it and the upstream *simOr67a.D* copy. We limited the total number of significant motifs to 12 for the comparative analysis.

## 6. Acknowledgements

We thank Margarida Cardoso Moreira, Lucia Prieto-Godino, Juan Antonio Sanchez Alcañiz, Laurent Keller, Manyuan Long, and member of the Arguello lab for comments on earlier versions of the manuscript. Research J.R.A.’s lab is supported by the Swiss National Science Foundation grants PP00P3_176956 and 310030_201188. T.O.A. was supported by a Human Frontier Science Program Long-Term Fellowship (LT000461/2015-L) and a Swiss National Science Foundation Ambizione Grant (PZ00P3 185743). Research in R.B.’s laboratory was supported by ERC Consolidator and Advanced Grants (615094 and 833548, respectively) and the Swiss National Science Foundation. We thank Mohammed Khallaf, Markus Knaden, Ales Svatos and Jerrit Weissflog at the Max Planck Institute for Chemical Ecology for the synthesized (*R*)-actinidine, and John Carlson and David Stern for sharing transgenic fly lines.

## 7. Supplementary Figures

**Sup. Fig. 1.**
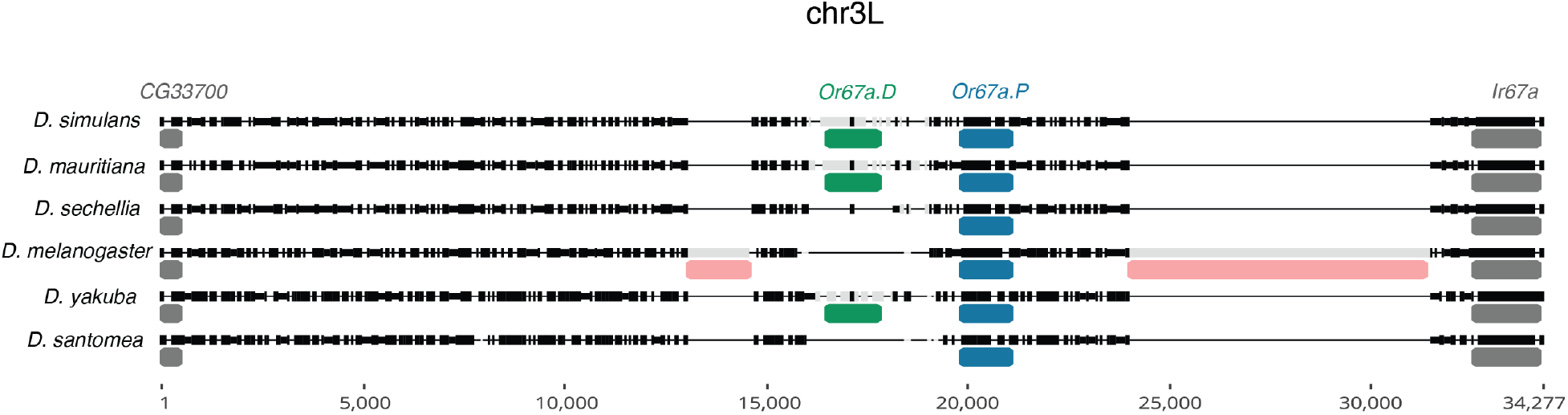
Alignment for the chromosome 3L interval containing *Or67a.D* and *Or67a.P* for six species. Higher sequence identity is indicated with black alignment blocks with low sequence identity indicated in grey. Thin horizontal lines are alignment gaps. Red annotations indicate locations of transposable elements. Chromosome position on the horizontal axis are relative to the extracted interval. See File S1 for the alignment in a flat file and File S3 for repeat annotations.

**Sup. Fig. 2.**
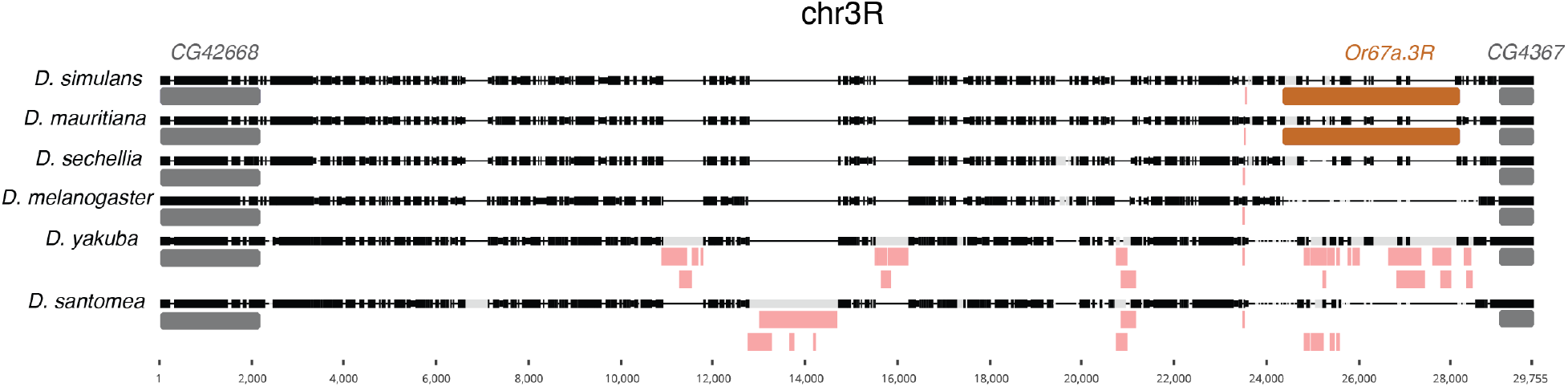
Alignment for the chromosome 3R interval containing *Or67a.D* and *Or67a.P* for six species. Higher sequence identity is indicated with black alignment blocks with low sequence identity indicated in grey. Thin horizontal lines are alignment gaps. Red annotations indicate locations of transposable elements. Chromosome position on the horizontal axis are relative to the extracted interval. See File S2 for the alignment in a flat file and File S3 for repeat annotations.

**Sup. Fig. 3.**
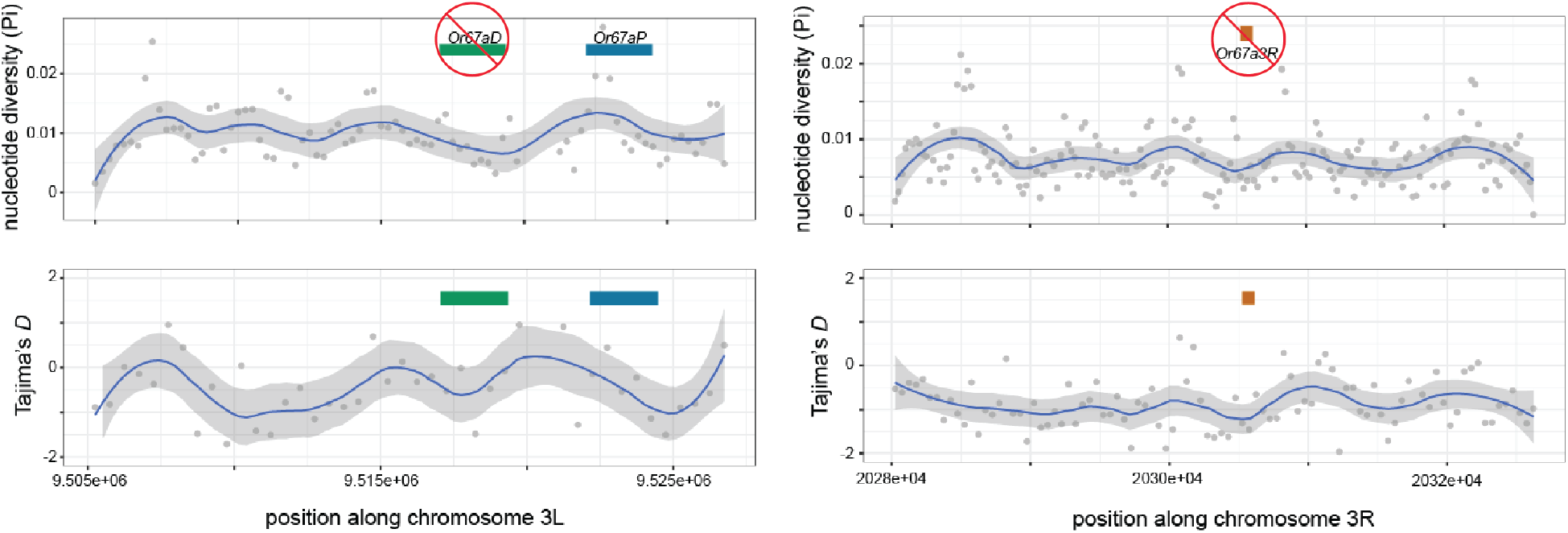
Nucleotide diversity and Tajima’s *D* over the over *D. melanogaster*’s chromosome regions containing the intact *Or67a.P* gene and the deleted *Or67a.D* and *Or67a.3R*. The regions containing the deleted *Or67a* paralogs do not show differences genetic diversity in comparison to the surrounding regions, as would be expected if the deletions were adaptive and swept in the population.

**Sup. Fig. 4.**
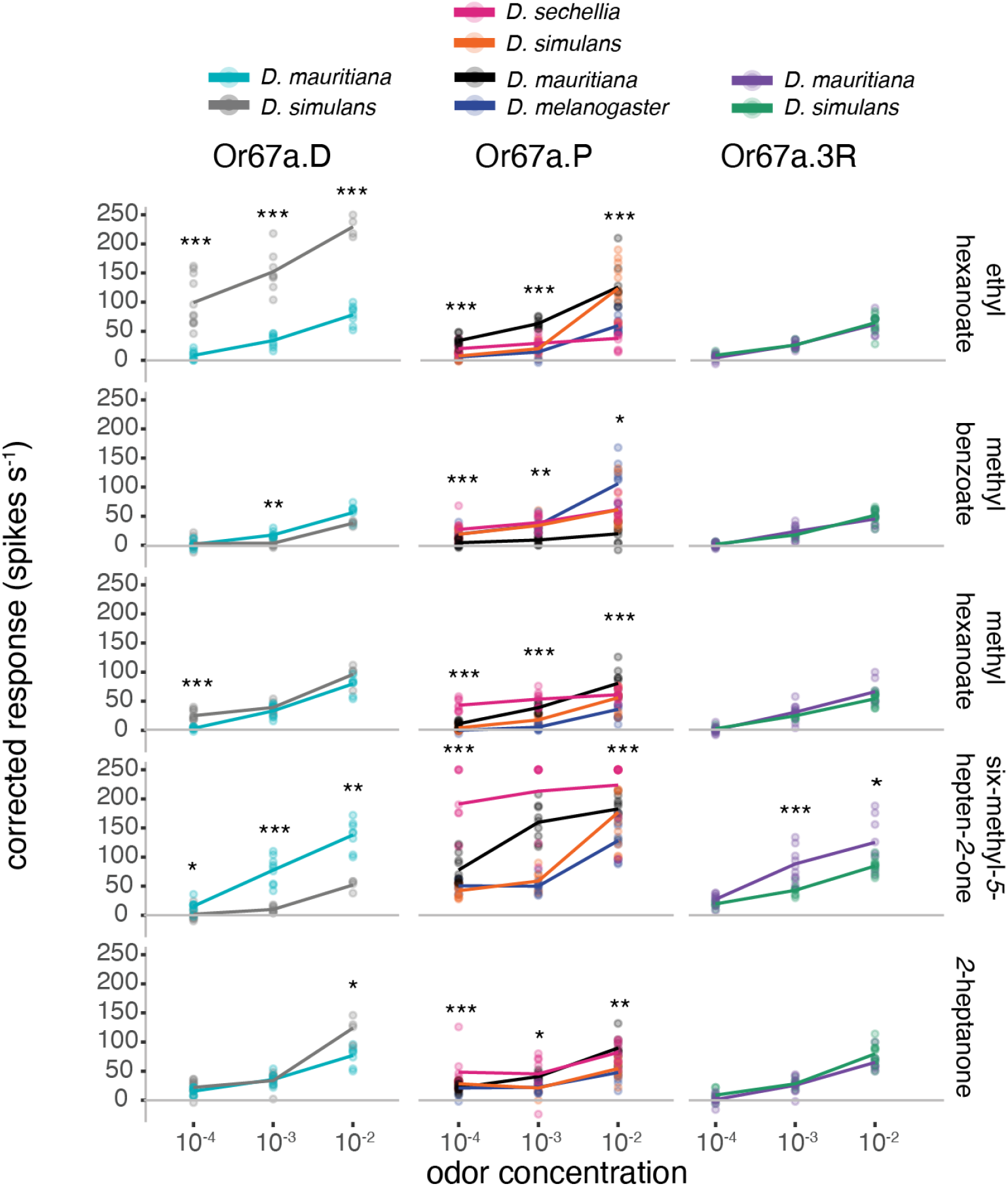
The full set of dose-response experiments for the subset of odors that evoked high or intermediate responses in our initial screen of nine odors (Fig. 2B). For simplicity, the level of significance indicated above each concentration’s comparison is only for the single comparison with the largest difference (see Table S4 for the full set of tests; \**p* < .05, \*\**p* < .01, \*\*\**p* < .001). Colors correspond to those in Fig. 2D. *p*-value correction for multiple comparisons was done using the Holm method. Sample sizes = 4-12).

**Sup. Fig. 5.**
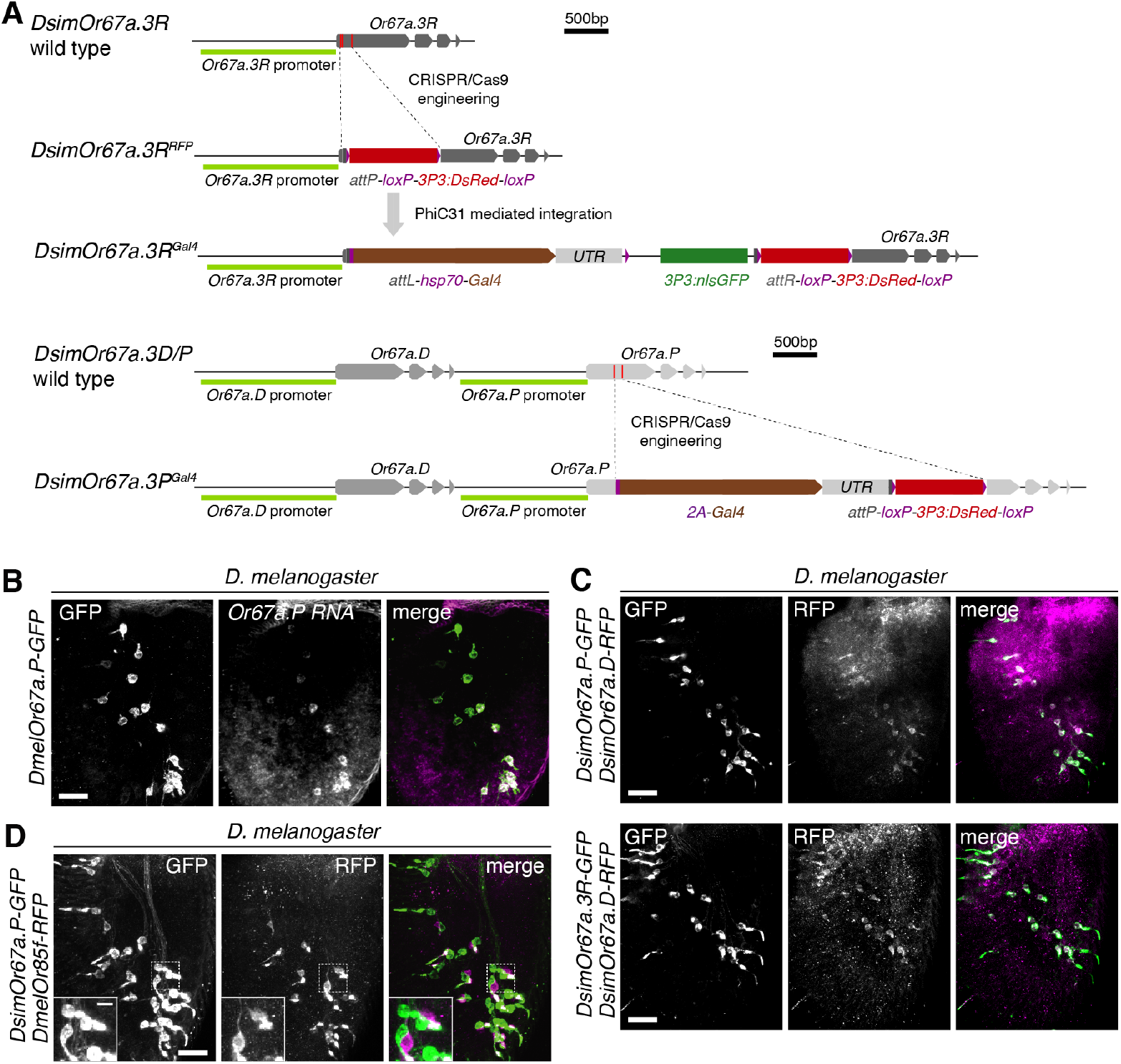
**(A)** Schematics of the wild-type and knock-in Gal4 transcriptional reporter alleles at the *DsimOr67a.3R* (top) and *DsimOr67a.D/P* (bottom) loci. The first was created via a two-step process (CRISPR/Cas9 engineering + PhiC31 mediated integration) while the latter was resulting from a direct CRISPR/Cas9 mediated insertion. **(B)** Antennal co-expression of the *DmelOr67a.P-GFP* transcriptional reporter and *Or67a.P* RNA in *D. melanogaster*. Scale bar = 25 *µ*m. **(C)** Antennal co-expression of the *simOr67a.P-GFP* and *simOr67a.D-RFP* transcriptional reporters (top) and the *simOr67a.3R-GFP* and *simOr67a.D-RFP* transcriptional reporters (bottom) in *D. melanogaster*. Scale bar = 25 *µ*m. **(D)** Pairing of the *simOr67a.P-GFP* and *melOr85f-RFP* transcriptional reporter in neighboring neurons in the antenna of *D. melanogaster*. Scale bar = 25 *µ*m. Inset scale bar = 5 *µ*m.

## 8. Supplemental Tables

- Table S1. Codeml table of models tested and likelihoods
- Table S2. Codeml table of likelihood ratio tests
- Table S3. Relative effects of odors on receptor responses based on the NPMV analysis
- Table S4. Dose-response tests
- Table S5. Pairwise identity between *Or67a* gene sequences
- Table S6. Pairwise identity between *Or67a* promoter sequences
- Table S7. Table with MEME results
- Table S8. Oligonucleotides used in this study

## 9. Supplemental Files

- File S1. Extending alignment for the 3L region containing *Or67a.D* and *Or67a.P*
- File S2. Extending alignment for the 3R region containing *Or67a.3R*
- File S3. RepeatMasker output used in Fig. S1 and S2.
- File S4. Codeml output files form models used in Table S1
- File S5. *D. simulans* and *D. melanogaster* polymorphism summaries
- File S6. Odor response data from electrophysiology experiments: 10−2
- File S7. Odor response data from electrophysiology experiments: dose-responses
- File S8. Odor response data from electrophysiology experiments: fluorescent-guided SSR
- File S9-11. *D. simulans* Sanger sequences generated in this study

## References

1. H. M. Robertson and K. W. Wanner, “The chemoreceptor superfamily in the honey bee, apis mellifera: expansion of the odorant, but not gustatory, receptor family,” Genome Res 16, 1395–403 (2006).

2. C. S. McBride and J. R. Arguello, “Five drosophila genomes reveal nonneutral evolution and the signature of host specialization in the chemoreceptor superfamily,” Genetics 177, 1395–416 (2007).

3. H. M. Robertson, “Molecular evolution of the major arthropod chemoreceptor gene families,” Annu. Rev Entomol 64, 227–242 (2019).

4. Y. Gilad, V. Wiebe, M. Przeworski, D. Lancet, and S. Pääbo, “Loss of olfactory receptor genes coincides with the acquisition of full trichromatic vision in primates,” PLoS Biol 2, E5 (2004).

5. G. M. Hughes, E. S. M. Boston, J. A. Finarelli, W. J. Murphy, D. G. Higgins, and E. C. Teeling, “The birth and death of olfactory receptor gene families in mammalian niche adaptation,” Mol Biol Evol 35, 1390–1406 (2018).

6. Y. Niimura, A. Matsui, and K. Touhara, “Extreme expansion of the olfactory receptor gene repertoire in african elephants and evolutionary dynamics of orthologous gene groups in 13 placental mammals,” Genome Res 24, 1485–96 (2014).

7. H. Robertson and K. Wanner, “The chemoreceptor superfamily in the honey bee, apis mellifera: Expansion of the odorant, but not gustatory, receptor family,” Genome Res. 16, 1395 (2006).

8. S. K. McKenzie, M. E. Winston, F. Grewe, G. Vargas Asensio, N. Rodríguez-Hernández, B. E. R. Rubin, C. Murillo-Cruz, C. von Beeren, C. S. Moreau, G. Suen, A.A. Pinto-Tomás, and D. J. C. Kronauer, “The genomic basis of army ant chemosensory adaptations,” Mol Ecol (2021).

9. H. Zhao, J. Li, and J. Zhang, “Molecular evidence for the loss of three basic tastes in penguins,” Curr Biol 25, R141–2 (2015).

10. H. Zhao, J.-R. Yang, H. Xu, and J. Zhang, “Pseudogenization of the umami taste receptor gene tas1r1 in the giant panda coincided with its dietary switch to bamboo,” Mol Biol Evol 27, 2669–73 (2010).

11. M. Nei and A. P. Rooney, “Concerted and birth-and-death evolution of multigene families,” Annu. Rev Genet. 39, 121–52 (2005).

12. S. Guo and J. Kim, “Molecular evolution of drosophila odorant receptor genes,” Mol Biol Evol 24, 1198–207 (2007).

13. Z. Zheng, J. S. Lauritzen, E. Perlman, C. G. Robinson, M. Nichols, D. Milkie, O. Torrens, J. Price, C. B. Fisher, N. Sharifi, S. A. Calle-Schuler, L. Kmecova, I. J. Ali, B. Karsh, E. T. Trautman, J. A. Bogovic, P. Hanslovsky, G. S. X. E. Jefferis, M. Kazhdan, K. Khairy, S. Saalfeld, R. D. Fetter, and D. D. Bock, “A complete electron microscopy volume of the brain of adult drosophila melanogaster,” Cell 174, 730–743.e22 (2018).

14. C. Eschbach, A. Fushiki, M. Winding, B. Afonso, I. V. Andrade, B. T. Cocanougher, K. Eichler, R. Gepner, G. Si, J. Valdes-Aleman, M. Gershow, G. S. Jefferis, J. W. Truman, R. D. Fetter, A. Samuel, A. Cardona, and M. Zlatic, “Circuits for integrating learnt and innate valences in the fly brain,” bioRxiv (2020).

15. A. M. Allen, M. C. Neville, S. Birtles, V. Croset, C. D. Treiber, S. Waddell, and S. F. Goodwin, “A single-cell transcriptomic atlas of the adult drosophila ventral nerve cord,” Elife 9 (2020).

16. H. Li, J. Janssens, M. De Waegeneer, S. S. Kolluru, K. Davie, V. Gardeux, W. Saelens, F. David, M. Brbić, J. Leskovec, C. N. McLaughlin, Q. Xie, R. C. Jones, K. Brueckner, J. Shim, S. G. Tattikota, F. Schnorrer, K. Rust, T. G. Nystul, Z. Carvalho-Santos, C. Ribeiro, S. Pal, T. M. Przytycka, A. M. Allen, S. F. Goodwin, C. W. Berry, M. T. Fuller, H. White-Cooper, E. L. Matunis, S. DiNardo, A. Galenza, L. E. O’Brien, J. A. T. Dow, F. Consortium, H. Jasper, B. Oliver, N. Perrimon, B. Deplancke, S. R. Quake, L. Luo, and S. Aerts, “Fly cell atlas: a single-cell transcriptomic atlas of the adult fruit fly,” bioRxiv (2021).

17. T. O. Auer, M. P. Shahandeh, and R. Benton, “Drosophila sechellia: A genetic model for behavioral evolution and neuroecology,” Annu. Rev Genet. (2021).

18. T. O. Auer, M. A. Khallaf, A. F. Silbering, G. Zappia, K. Ellis, R. Álvarez-Ocaña, J. R. Arguello, B. S. Hansson, G. S. X. E. Jefferis, S. J. C. Caron, M. Knaden, and R. Benton, “Olfactory receptor and circuit evolution promote host specialization,” Nature 579, 402–408 (2020).

19. L. L. Prieto-Godino, R. Rytz, S. Cruchet, B. Bargeton, L. Abuin, A. F. Silbering, V. Ruta, M. Dal Peraro, and R. Benton, “Evolution of acid-sensing olfactory circuits in drosophilids,” Neuron 93, 661–676.e6 (2017).

20. Y. Ding, J. L. Lillvis, J. Cande, G. J. Berman, B. J. Arthur, X. Long, M. Xu, B. J. Dickson, and D. L. Stern, “Neural evolution of context-dependent fly song,” Curr. Biol. 29, 1089–1099.e7 (2019).

21. L. F. Seeholzer, M. Seppo, D. L. Stern, and V. Ruta, “Evolution of a central neural circuit underlies drosophila mate preferences,” Nature 559, 564–569 (2018).

22. M. C. Larsson, A. I. Domingos, W. D. Jones, M. E. Chiappe, H. Amrein, and L. B. Vosshall, “Or83b encodes a broadly expressed odorant receptor essential for drosophila olfaction,” Neuron 43, 703–14 (2004).

23. S. Serizawa, K. Miyamichi, H. Nakatani, M. Suzuki, M. Saito, Y. Yoshihara, and H. Sakano, “Negative feedback regulation ensures the one receptor-one olfactory neuron rule in mouse,” Science 302, 2088–94 (2003).

24. J. W. Lewcock and R. R. Reed, “A feedback mechanism regulates monoallelic odorant receptor expression,” Proc Natl Acad Sci U S A 101, 1069–74 (2004).

25. A. Gardiner, D. Barker, R. K. Butlin, W. C. Jordan, and M. G. Ritchie, “Drosophila chemoreceptor gene evolution: selection, specialization and genome size,” Mol Ecol 17, 1648–57 (2008).

26. S. Ramasamy, L. Ometto, C. M. Crava, S. Revadi, R. Kaur, D. S. Horner, D. Pisani, T. Dekker, G. Anfora, and O. Rota-Stabelli, “The evolution of olfactory gene families in drosophila and the genomic basis of chemical-ecological adaptation in drosophila suzukii,” Genome Biol Evol 8, 2297–311 (2016).

27. D. J. Obbard, J. Maclennan, K.-W. Kim, A. Rambaut, P. M. O’Grady, and F. M. Jiggins, “Estimating divergence dates and substitution rates in the drosophila phylogeny,” Mol. Biol. Evol. 29, 3459–3473 (2012).

28. L. Ometto, A. Cestaro, S. Ramasamy, A. Grassi, S. Revadi, S. Siozios, M. Moretto, P. Fontana, C. Varotto, D. Pisani, T. Dekker, N. Wrobel, R. Viola, I. Pertot, D. Cavalieri, M. Blaxter, G. Anfora, and O. Rota-Stabelli, “Linking genomics and ecology to investigate the complex evolution of an invasive drosophila pest,” Genome Biol Evol 5, 745–57 (2013).

29. H. K. M. Dweck, S. A. M. Ebrahim, T. Retzke, V. Grabe, J. Weißflog, A. Svatoš, B. S. Hansson, and M. Knaden, “The olfactory logic behind fruit odor preferences in larval and adult drosophila,” Cell Reports 23, 2524–2531 (2018).

30. D. Garrigan, S. B. Kingan, A. J. Geneva, P. Andolfatto, A. G. Clark, K. R. Thornton, and D. C. Presgraves, “Genome sequencing reveals complex speciation in the drosophila simulans clade,” Genome Res 22, 1499–511 (2012).

31. D. R. Schrider, J. Ayroles, D. R. Matute, and A. D. Kern, “Supervised machine learning reveals introgressed loci in the genomes of drosophila simulans and d. sechellia,” PLOS Genet. 14, 1–29 (2018).

32. J. A. Butterwick, J. Del Mármol, K. H. Kim, M. A. Kahlson, J. A. Rogow, T. Walz, and V. Ruta, “Cryo-em structure of the insect olfactory receptor orco,” Nature 560, 447–452 (2018).

33. J. Del Mármol, M. A. Yedlin, and V. Ruta, “The structural basis of odorant recognition in insect olfactory receptors,” Nature (2021).

34. J. H. McDonald and M. Kreitman, “Adaptive protein evolution at the adh locus in drosophila,” Nature 351, 652–654 (1991).

35. N. G. C. Smith and A. Eyre-Walker, “Adaptive protein evolution in drosophila,” Nature 415, 1022–1024 (2002).

36. J. R. Arguello, M. Cardoso-Moreira, J. K. Grenier, S. Gottipati, A. G. Clark, and R. Benton, “Extensive local adaptation within the chemosensory system following drosophila melanogaster’s global expansion,” Nat Commun 7, ncomms11855 (2016).

37. S. Mansourian, A. Enjin, E. V. Jirle, V. Ramesh, G. Rehermann, P. G. Becher, J. E. Pool, and M. C. Stensmyr, “Wild african drosophila melanogaster are seasonal specialists on marula fruit,” Curr Biol 28, 3960–3968.e3 (2018).

38. R. Albalat and C. Cañestro, “Evolution by gene loss,” Nat Rev Genet. 17, 379–91 (2016).

39. F. Tajima, “Statistical Method for Testing the Neutral Mutation Hypothesis by DNA Polymorphism,” Genetics 123, 585–595 (1989).

40. P. A. Hohenlohe, P. C. Phillips, and W. A. Cresko, “Using population genomics to detect selection in natural populations: Key concepts and methodological considerations,” Int J Plant Sci 171, 1059–1071 (2010).

41. A. A. Dobritsa, W. van der Goes van Naters, C. G. Warr, R. A. Steinbrecht, and J. R. Carlson, “Integrating the molecular and cellular basis of odor coding in the drosophila antenna,” Neuron 37, 827–41 (2003).

42. E. Hallem and J. Carlson, “Coding of odors by a receptor repertoire,” Cell 125, 143–160 (2006).

43. E. Hallem, M. Ho, and J. Carlson, “The molecular basis of odor coding in the drosophila antenna,” Cell 117, 965–979 (2004).

44. C. G. Galizia, D. Münch, M. Strauch, A. Nissler, and S. Ma, “Integrating heterogeneous odor response data into a common response model: A door to the complete olfactome,” Chem Senses 35, 551–63 (2010).

45. W. W. Burchett, A. R. Ellis, S. W. Harrar, and A. C. Bathke, “Nonparametric inference for multivariate data: The r package npmv,” J. Stat. Softw. 76 (2017).

46. T. Dekker, I. Ibba, K. P. Siju, M. C. Stensmyr, and B. S. Hansson, “Olfactory shifts parallel superspecialism for toxic fruit in drosophila melanogaster sibling, d. sechellia,” Curr Biol 16, 101–9 (2006).

47. M. C. Stensmyr, T. Dekker, and B. S. Hansson, “Evolution of the olfactory code in the drosophila melanogaster subgroup,” Proc Biol Sci 270, 2333–40 (2003).

48. M. A. Khallaf, R. Cui, J. Weißflog, M. Erdogmus, A. Svatoš, H. K. M. Dweck, D. R. Valenzano, B. S. Hansson, and M. Knaden, “Large-scale characterization of sex pheromone communication systems in drosophila,” Nat Commun 12, 4165 (2021).

49. M. A. Khallaf, T. O. Auer, V. Grabe, A. Depetris-Chauvin, B. Ammagarahalli, D.-D. Zhang, S. Lavista-Llanos, F. Kaftan, J. Weißflog, L. M. Matzkin, S. M. Rollmann, C. Löfstedt, A. Svatoš, H. K. M. Dweck, S. Sachse, R. Benton, B. S. Hansson, and M. Knaden, “Mate discrimination among subspecies through a conserved olfactory pathway,” Sci Adv 6, eaba5279 (2020).

50. M. C. Stensmyr, H. K. M. Dweck, A. Farhan, I. Ibba, A. Strutz, L. Mukunda, J. Linz, V. Grabe, K. Steck, S. Lavista-Llanos, D. Wicher, S. Sachse, M. Knaden, P. Becher, Y. Seki, and B. S. Hansson, “A conserved dedicated olfactory circuit for detecting harmful microbes in drosophila,” Cell 151, 1345–1357 (2012).

51. L. B. Vosshall and R. F. Stocker, “Molecular architecture of smell and taste in drosophila,” Annu. Rev. Neurosci. 30, 505–533 (2007).

52. A. Ray, W. v. d. G. van Naters, and J. R. Carlson, “A regulatory code for neuron-specific odor receptor expression,” PLoS Biol 6, e125 (2008).

53. A. Ray, W. v. d. G. van Naters, T. Shiraiwa, and J. R. Carlson, “Mechanisms of odor receptor gene choice in drosophila,” Neuron 53, 353–69 (2007).

54. A. L. Goldman, W. Van der Goes van Naters, D. Lessing, C. G. Warr, and J. R. Carlson, “Coexpression of two functional odor receptors in one neuron,” Neuron 45, 661–6 (2005).

55. M. Aguadé, “Nucleotide and copy-number polymorphism at the odorant receptor genes or22a and or22b in drosophila melanogaster,” Mol Biol Evol 26, 61–70 (2009).

56. S. Lebreton, F. Borrero-Echeverry, F. Gonzalez, M. Solum, E. A. Wallin, E. Hedenström, B. S. Hansson, A.-L. Gustavsson, M. Bengtsson, G. Birgersson, W. B. Walker, 3rd, H. K. M. Dweck, P. G. Becher, and P. Witzgall, “A drosophila female pheromone elicits species-specific long-range attraction via an olfactory channel with dual specificity for sex and food,” BMC Biol 15, 88 (2017).

57. M. de Bruyne, R. Smart, E. Zammit, and C. G. Warr, “Functional and molecular evolution of olfactory neurons and receptors for aliphatic esters across the drosophila genus,” J Comp Physiol A Neuroethol Sens Neural Behav Physiol 196, 97–109 (2010).

58. K. Scott, “Gustatory processing in drosophila melanogaster,” Annu. Rev. Entomol. 63, 15–30 (2018).

59. A. Dahanukar, Y.-T. Lei, J. Y. Kwon, and J. R. Carlson, “Two gr genes underlie sugar reception in drosophila,” Neuron 56, 503–16 (2007).

60. S. Fujii, A. Yavuz, J. Slone, C. Jagge, X. Song, and H. Amrein, “Drosophila sugar receptors in sweet taste perception, olfaction, and internal nutrient sensing,” Curr. Biol. 25, 621–627 (2015).

61. Y. Jiao, S. J. Moon, X. Wang, Q. Ren, and C. Montell, “Gr64f is required in combination with other gustatory receptors for sugar detection in drosophila,” Curr Biol 18, 1797–801 (2008).

62. J. Y. Kwon, A. Dahanukar, L. A. Weiss, and J. R. Carlson, “Molecular and cellular organization of the taste system in the drosophila larva,” J Neurosci 31, 15300–9 (2011).

63. J. Slone, J. Daniels, and H. Amrein, “Sugar receptors in drosophila,” Curr Biol 17, 1809–16 (2007).

64. H. K. M. Dweck and J. R. Carlson, “Molecular logic and evolution of bitter taste in drosophila,” Curr Biol 30, 17–30.e3 (2020).

65. A. Ganguly, L. Pang, V.-K. Duong, A. Lee, H. Schoniger, E. Varady, and A. Dahanukar, “A molecular and cellular context-dependent role for ir76b in detection of amino acid taste,” Cell Rep 18, 737–750 (2017).

66. J. M. Tauber, E. B. Brown, Y. Li, M. E. Yurgel, P. Masek, and A. C. Keene, “A subset of sweet-sensing neurons identified by ir56d are necessary and sufficient for fatty acid taste,” PLoS Genet. 13, e1007059 (2017).

67. F. Port and S. L. Bullock, “Augmenting crispr applications in drosophila with trna-flanked sgrnas,” Nat Methods 13, 852–4 (2016).

68. F. Port, H.-M. Chen, T. Lee, and S. L. Bullock, “Optimized crispr/cas tools for efficient germline and somatic genome engineering in drosophila,” Proc Natl Acad Sci U S A 111, E2967–76 (2014).

69. S. J. Gratz, F. P. Ukken, C. D. Rubinstein, G. Thiede, L. K. Donohue, A. M. Cummings, and K.M. O’Connor-Giles, “Highly specific and efficient crispr/cas9-catalyzed homology-directed repair in drosophila,” Genetics 196, 961–71 (2014).

70. S. Kondo, T. Takahashi, N. Yamagata, Y. Imanishi, H. Katow, S. Hiramatsu, K. Lynn, A. Abe, A. Kumaraswamy, and H. Tanimoto, “Neurochemica organization of the drosophila brain visualized by endogenously tagged neurotransmitter receptors,” Cell Rep 30, 284–297.e5 (2020).

71. J. Bischof, R. K. Maeda, M. Hediger, F. Karch, and K. Basler, “An optimized transgenesis system for drosophila using germ-line-specific phic31 integrases,” Proc Natl Acad Sci U S A 104, 3312–7 (2007).

72. C. Han, L. Y. Jan, and Y.-N. Jan, “Enhancer-driven membrane markers for analysis of nonautonomous mechanisms reveal neuron-glia interactions in drosophila,” Proc Natl Acad Sci U S A 108, 9673–8 (2011).

73. D. M. Gohl, M. A. Silies, X. J. Gao, S. Bhalerao, F. J. Luongo, C.-C. Lin, C. J. Potter, and T. R. Clandinin, “A versatile in vivo system for directed dissection of gene expression patterns,” Nat Methods 8, 231–7 (2011).

74. R. Benton and A. Dahanukar, “Electrophysiological recording from drosophila olfactory sensilla,” Cold Spring Harb Protoc 2011, 824–38 (2011).

75. S. A. M. Ebrahim, H. K. M. Dweck, J. Stökl, J. E. Hofferberth, F. Trona, K. Weniger, J. Rybak, Y. Seki, M. C. Stensmyr, S. Sachse, B. S. Hansson, and M. Knaden, “Drosophila avoids parasitoids by sensing their semiochemicals via a dedicated olfactory circuit,” PLoS Biol 13, e1002318 (2015).

76. C.-C. Lin and C. J. Potter, “Re-classification of drosophila melanogaster trichoid and intermediate sensilla using fluorescence-guided single sensillum recording,” PLoS One 10, e0139675 (2015).

77. R. C. Team, “R: A language and environment for statistical computing,” (2020).

78. H. Wickham, ggplot2: Elegant Graphics for Data Analysis (Springer-Verlag New York, 2016).

79. U. Ligges and M. Mächler, “Scatterplot3d - an r package for visualizing multivariate data,” J. Stat. Softw. 8, 1–20 (2003).

80. D. J. Stekhoven and P. Bühlmann, “Missforest–non-parametric missing value imputation for mixed-type data,” Bioinformatics 28, 112–8 (2012).

81. M. Saina and R. Benton, “Visualizing olfactory receptor expression and localization in drosophila,” Methods Mol Biol 1003, 211–28 (2013).

82. A. F. Silbering, R. Rytz, Y. Grosjean, L. Abuin, P. Ramdya, G. S. X. E. Jefferis, and R. Benton, “Complementary function and integrated wiring of the evolutionarily distinct drosophila olfactory subsystems,” J Neurosci 31, 13357–75 (2011).

83. J.A. Sánchez-Alcañiz, G. Zappia, F. Marion-Poll, and R. Benton, “A mechanosensory receptor required for food texture detection in drosophila,” Nat Commun 8, 14192 (2017).

84. J. Schindelin, I. Arganda-Carreras, E. Frise, V. Kaynig, M. Longair, T. Pietzsch, S. Preibisch, C. Rueden, S. Saalfeld, B. Schmid, J.-Y. Tinevez, D. J. White, V. Hartenstein, K. Eliceiri, P. Tomancak, and A. Cardona, “Fiji: an open-source platform for biological-image analysis,” Nat Methods 9, 676–82 (2012).

85. F. Sievers, A. Wilm, D. Dineen, T. J. Gibson, K. Karplus, W. Li, R. Lopez, H. McWilliam, M. Remmert, J. Söding, J. D. Thompson, and D. G. Higgins, “Fast, scalable generation of high-quality protein multiple sequence alignments using clustal omega,” Mol. Syst. Biol. 7, 539 (2011).

86. F. Ronquist and J. P. Huelsenbeck, “Mrbayes 3: Bayesian phylogenetic inference under mixed models,” Bioinformatics 19, 1572–4 (2003).

87. Z. Yang, “Paml 4: phylogenetic analysis by maximum likelihood,” Mol Biol Evol 24, 1586–91 (2007).

88. B. Xu and Z. Yang, “Pamlx: a graphical user interface for paml,” Mol Biol Evol 30, 2723–4 (2013).

89. S. A. Signor, F. N. New, and S. Nuzhdin, “A large panel of drosophila simulans reveals an abundance of common variants,” Genome Biol Evol 10, 189–206 (2018).

90. P. Danecek, A. Auton, G. Abecasis, C. A. Albers, E. Banks, M. A. DePristo, R. E. Handsaker, G. Lunter, G. T. Marth, S. T. Sherry, G. McVean, R. Durbin, and. G. P. A. Group, “The variant call format and vcftools,” Bioinformatics 27, 2156–2158 (2011).

91. J. K. Grenier, J. R. Arguello, M. Cardoso Moreira, S. Gottipati, J. Mohammed, S. R. Hackett, R. Boughton, A. J. Greenberg, and A. G. Clark, “Global diversity lines - a five-continent reference panel of sequenced drosophila melanogaster strains,” G3 pp. 593–603 (2015).

92. M. Chakraborty, C.-H. Chang, D. E. Khost, J. Vedanayagam, J. R. Adrion, Y. Liao, K. L. Montooth, C. D. Meiklejohn, A. M. Larracuente, and J. J. Emerson, “Evolution of genome structure in the drosophila simulans species complex,” Genome Res 31, 380–396 (2021).

93. D. E. Miller, C. Staber, J. Zeitlinger, and R. S. Hawley, “Highly contiguous genome assemblies of 15 drosophila species generated using nanopore sequencing,” G3 (Bethesda) 8, 3131–3141 (2018).

94. A. Smit, R. Hubley, and P. Green, “Repeatmasker open-4.0,”.

95. T. L. Bailey, N. Williams, C. Misleh, and W. W. Li, “Meme: discovering and analyzing dna and protein sequence motifs,” Nucleic Acids Res. 34, W369–W373 (2006).

96. T. L. Bailey, M. Boden, F. A. Buske, M. Frith, C. E. Grant, L. Clementi, J. Ren, W. W. Li, and W. S. Noble, “Meme suite: tools for motif discovery and searching,” Nucleic Acids Res 37, W202–8 (2009).

