## Supplementary material for "Odorant receptor copy number change, co-expression, and positive selection establish peripheral coding differences between fly species": sup. file 3: Dmel_3R_region.html

[ close all hsps ]
[ open all hsps ]

```
                                          position in query-                           -position in repeat-
          %    %    %    query                               C matching  repeat        (left)  end   begin  linkage
+ score  div. del. ins.  sequence         begin  end  (left) + repeat    class/family  begin   end   (left) id/graphic
```

```
+  236   11.8  0.0  0.0  UnnamedSequence    281   314 (1032) + LINEJ1_DM LINE/I-Jockey   4469   4502  (517)   1
```

```
ANNOTATION EVIDENCE:
  236  11.76 0.00 0.00  UnnamedSequence    281   314   1032 + LINEJ1_DM LINE/I-Jockey   4469   4502    517    
236 11.76 0.00 0.00 UnnamedSequence 281 314 (1032) LINEJ1_DM#LINE/I-Jockey 4469 4502 (517) m_b1s001i0

  UnnamedSequen        281 TGTTCTACGGCCTTCAGGTATACAGTATTTCTGC 314
                                        v         i i   v    
  LINEJ1_DM#LIN       4469 TGTTCTACGGCCTGCAGGTATACGGCATTGCTGC 4502

Matrix = 20p43g.matrix
Kimura (with divCpGMod) = 9.72
Transitions / transversions = 1.00 (2/2)
Gap_init rate = 0.00 (0 / 33), avg. gap size = 0.0 (0 / 0)
```
