## Supplementary material for "Odorant receptor copy number change, co-expression, and positive selection establish peripheral coding differences between fly species": sup. file 3: RM2sequpload_1628786844.out.html

[ close all hsps ]
[ open all hsps ]

```
                                          position in query-                            -position in repeat-
          %    %    %    query                               C matching repeat          (left)  end   begin  linkage
+ score  div. del. ins.  sequence         begin  end  (left) + repeat   class/family    begin   end   (left) id/graphic
```

```
+  779   24.3  6.1  9.7  UnnamedSequence     11   534 (1431) + Transib5 DNA/CMC-Transib      1    507 (2494)   1
```

```
ANNOTATION EVIDENCE:
  779  24.30 6.11 9.66  UnnamedSequence     11   534   1431 + Transib5 DNA/CMC-Transib      1    507   2494    
779 24.30 6.11 9.66 UnnamedSequence 11 534 (1431) Transib5#DNA/CMC-Transib 1 507 (2494) m_b1s001i0

  UnnamedSequen         11 CACAGTGGTTGCGCCCATAGGCCAAAAATGAAAAAAAAATGTTTCCAAAT 60
                                     v  i - v           -        ?i  i vv v  
  Transib5#DNA/          1 CACAGTGGTTCCGTC-AGAGGCCAAAAAT-AAAAAAAANCGTCTAAATAT 48

  UnnamedSequen         61 AATTACGAA--AAGTAACCAGATTGTCTCAAAAATGCCCAAATATGTGTA 108
                            v     v --         i        v   vv i        v vi 
  Transib5#DNA/         49 ATTTACGCATAAAGTAACCAAATTGTCTCCAAATAGTCCAAATATTTCCA 98

  UnnamedSequen        109 TGTCAAAACC-GTTTTGACTT-AACTCTAACGGGACATTCTAAAAATATA 156
                           - ?       -     vi   -  vi       vv v           ? 
  Transib5#DNA/         99 -GNCAAAACCAGTTTTTGCTTCAAACCTAACGGTTCTTTCTAAAAATANA 147

  UnnamedSequen        157 GTGTAAAGGGA-GAACAACTGTTCATTTTTGCAGCACA---GCG-GAGTT 201
                           i -     vv - v   vv         v  ?   iv ---  i- vi  
  Transib5#DNA/        148 AT-TAAAGTTATGCACACGTGTTCATTTGTGNAGCGAAATTGCACGCATT 196

  UnnamedSequen        202 TTAATTGCTTTTCGTCGCA--ACAAAAGTGAATTTCAAAGTGGTCAAACG 249
                           v     v  -------   --v      ii  i       vvi   v  v
  Transib5#DNA/        197 GTAATTTCT-------GCATTCCAAAAGCAAACTTCAAAGGCATCATACT 239

  UnnamedSequen        250 TGCGATTGATAGGTCCAATTTTAATTATTTAAAATGAATTTTAAAGTTTC 299
                            v ii viiv  v i i    v   v          i          vvv
  Transib5#DNA/        240 TTCAGTGAGGAGCTTCGATTTAAATAATTTAAAATGGATTTTAAAGTAGA 289

  UnnamedSequen        300 AGGTATTTACTTTA---ATAAATTCTTATTGTGAAAT------TCATTGA 340
                                        i--- i   vv     ----    ------   v v 
  Transib5#DNA/        290 AGGTATTTACTTTGTTTACAAAAACTTAT----AAATAACTAATCAGTTA 335

  UnnamedSequen        341 ATTGATCTA--------TCATATTT---GGTGAATTTTTGATGAATTATA 379
                            i i vi  -------- v    i ---ii  ivi    v  ------ i
  Transib5#DNA/        336 ACTAAGTTAAGTAAAATTAATATCTTCAAATGGCCTTTTTAT------TG 379

  UnnamedSequen        380 CCTTTTTTGTGGACGTCATTTTCCTAGTTTAAGTAAACTTTTAGATATAT 429
                           i           --  -      v -     vv   i       v  ?i 
  Transib5#DNA/        380 TCTTTTTTGTGG--GT-ATTTTCAT-GTTTACTTAAGCTTTTAGTTANGT 425

  UnnamedSequen        430 TGCCCCCCGATCCCCCCTTAATGAGTGCTATCACTGCAATT-GTCAGTGT 478
                            --------------      ------v   i ----    -iv  vv  
  Transib5#DNA/        426 T--------------CCTTAA------ATATTA----AATTAAGCATAGT 451

  UnnamedSequen        479 TTTAAGGAAATAAAAATCGTAAGTCTAGCTGCAAATATTTGCAAACATAC 528
                               v iv       i iv i   ii      ii v i    ii ii i 
  Transib5#DNA/        452 TTTACGATAATAAAAGTTTTGAGTTCAGCTGCGGAAACTTGCGGATGTGC 501

  UnnamedSequen        529 GTGTAA 534
                           v     
  Transib5#DNA/        502 TTGTAA 507

Matrix = 20p43g.matrix
Kimura (with divCpGMod) = 28.30
Transitions / transversions = 0.85 (53/62)
Gap_init rate = 0.12 (61 / 523), avg. gap size = 1.33 (81 / 61)
```

```
+  240   27.0 11.2  0.0  UnnamedSequence    920  1008  (957) + Transib5 DNA/CMC-Transib   2307   2405  (596)   2
```

```
ANNOTATION EVIDENCE:
  240  26.97 11.24 0.00  UnnamedSequence    920  1008    957 + Transib5 DNA/CMC-Transib   2307   2405    596    
240 26.97 11.24 0.00 UnnamedSequence 920 1008 (957) Transib5#DNA/CMC-Transib 2307 2405 (596) m_b1s001i1

  UnnamedSequen        920 ATACACGGGCCTGCAATAATTGAAAATGCTCTCATATCGATCGGCGAGTT 969
                             v  i  ii     i    vv iv    vi vi i  i  i  v  ii 
  Transib5#DNA/       2307 ATTCATGGATCTGCAGTAATACAGCATGCGTTAGTGTCAATTGGAGAACT 2356

  UnnamedSequen        970 GTCAGAGGAAGCTGCTGAAT----CGATAC------AAAAAAATTTCGA 1008
                           v  i           v    ---- i    ------        ? v  
  Transib5#DNA/       2357 TTCGGAGGAAGCTGCGGAATCTAACAATACGGACTTAAAAAAATNTAGA 2405

Matrix = 20p43g.matrix
Kimura (with divCpGMod) = 32.94
Transitions / transversions = 1.40 (14/10)
Gap_init rate = 0.02 (2 / 88), avg. gap size = 5.00 (10 / 2)
```

```
+  261   15.6  0.0  2.2  UnnamedSequence   1438  1483  (482) + Transib5 DNA/CMC-Transib   2957   3001    (0)   3
```

```
ANNOTATION EVIDENCE:
  261  15.57 0.00 2.22  UnnamedSequence   1438  1483    482 + Transib5 DNA/CMC-Transib   2957   3001      0    
261 15.57 0.00 2.22 UnnamedSequence 1438 1483 (482) Transib5#DNA/CMC-Transib 2957 3001 (0) m_b1s001i2

  UnnamedSequen       1438 TAGAAGTATTTTTGGCCGCTCTCTCCATATAGGCGCAACCACTGTG 1483
                            i         v        -v       i i   v i        
  Transib5#DNA/       2957 TGGAAGTATTTATGGCCGCT-GCTCCATACAAGCGGAGCCACTGTG 3001

Matrix = 20p43g.matrix
Kimura (with divCpGMod) = 17.59
Transitions / transversions = 1.33 (4/3)
Gap_init rate = 0.02 (1 / 45), avg. gap size = 1.00 (1 / 1)
```

```
+  828   21.4  4.0  4.3  UnnamedSequence   1662  1964    (1) C Transib5 DNA/CMC-Transib (2699)    302      1   4
```

```
ANNOTATION EVIDENCE:
  828  21.38 3.96 4.30  UnnamedSequence   1662  1964      1 C Transib5 DNA/CMC-Transib      1    302   2699    
828 21.38 3.96 4.30 UnnamedSequence 1662 1964 (1) C Transib5#DNA/CMC-Transib (2699) 302 1 m_b1s001i3

  UnnamedSequen       1662 AAAGTAAATACCTGAAACTTTAAAATTCATTTTAAATAATTAAAATTGGA 1711
                                        vvv          i          v   v    i i 
C Transib5#DNA/        302 AAAGTAAATACCTTCTACTTTAAAATCCATTTTAAATTATTTAAATCGAA 253

  UnnamedSequen       1712 CCTATCAATCGCACGTTTGACCACTTTGAAATTCACTTTTGT--TGCGAC 1759
                           v  viiv ii v v  v   ivv       i  ii      v--   ---
C Transib5#DNA/        252 GCTCCTCACTGAAAGTATGATGCCTTTGAAGTTTGCTTTTGGAATGC--- 206

  UnnamedSequen       1760 GAAAAGCAATTAAAACTC-CGC---TGTGCTGCAAAAATGAACAGTTGTT 1805
                           ----  v     v  iv -i  --- vi   ?  v         vv   v
C Transib5#DNA/        205 ----AGAAATTACAATGCGTGCAATTTCGCTNCACAAATGAACACGTGTG 160

  UnnamedSequen       1806 CTC--CTTTACACTATATTTTTAGAATGTCCCGTTAGAGTT-AAGTCAAA 1852
                            vi--     - i ?           v vv       iv  -   iv   
C Transib5#DNA/        159 CATAACTTTA-ATTNTATTTTTAGAAAGAACCGTTAGGTTTGAAGCAAAA 111

  UnnamedSequen       1853 AC-GGTTTTGCCATACACATATTTGAGCATTTTTGAGACAATCTGGTTAC 1901
                             -       ? - iv v       ii vv   v        i       
C Transib5#DNA/        110 ACTGGTTTTGNC-TGGAAATATTTGGACTATTTGGAGACAATTTGGTTAC 62

  UnnamedSequen       1902 TT--TTCGTAAATATTTGGAAACGTTTTTTTTTTTCATTTTTGGCCTATG 1949
                             -- v         v vv i   ?        ---           v -
C Transib5#DNA/         61 TTTATGCGTAAATATATTTAGACGNTTTTTTTT---ATTTTTGGCCTCT- 16

  UnnamedSequen       1950 GACGCAACCACTGTG 1964
                               v          
C Transib5#DNA/         15 GACGGAACCACTGTG 1

Matrix = 20p43g.matrix
Kimura (with divCpGMod) = 24.71
Transitions / transversions = 0.59 (23/39)
Gap_init rate = 0.07 (20 / 302), avg. gap size = 1.25 (25 / 20)
```
