## Supplementary material for "Odorant receptor copy number change, co-expression, and positive selection establish peripheral coding differences between fly species": sup. file 3: RM2sequpload_1628860266.out.html

[ close all hsps ]
[ open all hsps ]

```
                                          position in query-                             -position in repeat-
          %    %    %    query                               C matching    repeat        (left)  end   begin  linkage
+ score  div. del. ins.  sequence         begin  end  (left) + repeat      class/family  begin   end   (left) id/graphic
```

```
+ 3048    5.0  6.5  0.2  UnnamedSequence     57   194  (299) + DNAREP1_DM  RC/Helitron        1    149  (445)   1
```

```
ANNOTATION EVIDENCE:
 3048   5.02 6.52 0.24  UnnamedSequence     57   194    299 + DNAREP1_DM  RC/Helitron        1    149    445    
3048 5.02 6.52 0.24 UnnamedSequence 57 194 (299) DNAREP1_DM#RC/Helitron 1 149 (445) m_b1s001i0

  UnnamedSequen         57 TTATACCCGTTACTCGTAGAGTAAAAGG-TATACTAAATTCGTTGAAAAG 105
                                                       -       i             
  DNAREP1_DM#RC          1 TTATACCCGTTACTCGTAGAGTAAAAGGGTATACTAGATTCGTTGAAAAG 50

  UnnamedSequen        106 TATGTAACAGGCAGAAGGAAGCGTTTCCGACCATATAAAGTATATATG-- 153
                                                                        ?i --
  DNAREP1_DM#RC         51 TATGTAACAGGCAGAAGGAAGCGTTTCCGACCATATAAAGTATATNCGCG 100

  UnnamedSequen        154 --------TATATTCTTGATCAGGATCAGTAGCCGAGTCGATCTGGCCA 194
                           --------                    i                    
  DNAREP1_DM#RC        101 CGCGCGCGTATATTCTTGATCAGGATCAATAGCCGAGTCGATCTGGCCA 149

Matrix = 20p43g.matrix
Kimura (with divCpGMod) = 5.04
Transitions / transversions = 1.00 (3/0)
Gap_init rate = 0.02 (3 / 137), avg. gap size = 3.67 (11 / 3)
```

```
+   32    5.4  0.0  0.0  UnnamedSequence    195   233  (260) + (TGTCCGTC)n Simple_repeat      1     39    (0)   2
```

```
ANNOTATION EVIDENCE:
   32   5.41 0.00 0.00  UnnamedSequence    195   233    260 + (TGTCCGTC)n Simple_repeat      1     39      0    
32 5.41 0.00 0.00 UnnamedSequence 195 233 (260) (TGTCCGTC)n#Simple_repeat 1 39 (0) c_b1s251i0

  UnnamedSequen        195 TGTCCGTCTGTCCGTCCGTCTGTCTGTCCGTCTGTCCGT 233
                                           i   i                  
  (TGTCCGTC)n#S          1 TGTCCGTCTGTCCGTCTGTCCGTCTGTCCGTCTGTCCGT 39

Matrix = Unknown
Transitions / transversions = 1.00 (2/0)
Gap_init rate = 0.00 (0 / 38), avg. gap size = 0.0 (0 / 0)
```

```
+ 3048    5.0  6.5  0.2  UnnamedSequence    234   493    (0) + DNAREP1_DM  RC/Helitron      150    423  (171)   1
```

```
ANNOTATION EVIDENCE:
 3048   5.02 6.52 0.24  UnnamedSequence    234   493      0 + DNAREP1_DM  RC/Helitron      150    423    171    
3048 5.02 6.52 0.24 UnnamedSequence 234 493 (0) DNAREP1_DM#RC/Helitron 150 423 (171) m_b1s001i0

  UnnamedSequen        234 --------------ATGAACGTCGAGATCTCAGGAACTACAAAAGCCAGA 269
                           --------------                         i      i   
  DNAREP1_DM#RC        150 GTCCGTCTGTCCGTATGAACGTCGAGATCTCAGGAACTATAAAAGCTAGA 199

  UnnamedSequen        270 AAGTTGAGATTAAGTATACAGACTCCAGGGACATAGACGCAGCGCAAGTT 319
                            i         ?  i       i  i  i                     
  DNAREP1_DM#RC        200 AGGTTGAGATTNAGCATACAGATTCTAGAGACATAGACGCAGCGCAAGTT 249

  UnnamedSequen        320 TGTCGATTCATGTTGCCACGCCCACTCTAACGCCCACAAACCGCCCAAAA 369
                              i  ii                                          
  DNAREP1_DM#RC        250 TGTTGACCCATGTTGCCACGCCCACTCTAACGCCCACAAACCGCCCAAAA 299

  UnnamedSequen        370 CTGCCACGCCCACACTTTTGAAAAA-TGTTTTGATATATTTTCATTTTTG 418
                                                    - v         -       v?  v
  DNAREP1_DM#RC        300 CTGCCACGCCCACACTTTTGAAAAANTTTTTTGATAT-TTTTCATANTTT 348

  UnnamedSequen        419 TATTGGTCTTGTAAATTTCTATCGATTTGCCAAAAAACTTTTTGCCACGC 468
                               i                                             
  DNAREP1_DM#RC        349 TATTAGTCTTGTAAATTTCTATCGATTTGCCAAAAAACTTTTTGCCACGC 398

  UnnamedSequen        469 CCACTCTAACGCCCACAAACCGCCC 493
                                         vv  i      
  DNAREP1_DM#RC        399 CCACTCTAACGCCCTAAAGCCGCCC 423

Matrix = 20p43g.matrix
Kimura (with divCpGMod) = 5.04
Transitions / transversions = 2.40 (12/5)
Gap_init rate = 0.01 (3 / 259), avg. gap size = 5.33 (16 / 3)
```
