## Supplementary material for "Odorant receptor copy number change, co-expression, and positive selection establish peripheral coding differences between fly species": sup. file 3: RM2sequpload_1628860748.out.html

[ close all hsps ]
[ open all hsps ]

```
                                          position in query-                          -position in repeat-
          %    %    %    query                               C matching   repeat      (left)  end   begin  linkage
+ score  div. del. ins.  sequence         begin  end  (left) + repeat     class/famil begin   end   (left) id/graphic
```

```
+  811    9.4  1.6  1.6  UnnamedSequence      4   132   (89) + DNAREP1_DM RC/Helitron    303    431  (163)   1
```

```
ANNOTATION EVIDENCE:
  811   9.45 1.55 1.55  UnnamedSequence      4   132     89 + DNAREP1_DM RC/Helitron    303    431    163    
811 9.45 1.55 1.55 UnnamedSequence 4 132 (89) DNAREP1_DM#RC/Helitron 303 431 (163) m_b1s001i0

  UnnamedSequen          4 GCTACGCCCACACTTTTGAAAAA-TGTTTTGAAATTTTTTA-ATTTTTGT 51
                             i                    - v      v      i - ?   -- 
  DNAREP1_DM#RC        303 GCCACGCCCACACTTTTGAAAAANTTTTTTGATATTTTTCATANTTT--T 350

  UnnamedSequen         52 ATTGGTCTTGTAAATTTCTATCGATTTGCCAAAAAACTTTCTGCCACGCC 101
                              i                                    i         
  DNAREP1_DM#RC        351 ATTAGTCTTGTAAATTTCTATCGATTTGCCAAAAAACTTTTTGCCACGCC 400

  UnnamedSequen        102 CACTATAACGCCTACAAACCGCCAAAAACTG 132
                               v       ivv  i     v       
  DNAREP1_DM#RC        401 CACTCTAACGCCCTAAAGCCGCCCAAAACTG 431

Matrix = 20p43g.matrix
Kimura (with divCpGMod) = 10.21
Transitions / transversions = 1.00 (6/6)
Gap_init rate = 0.03 (4 / 128), avg. gap size = 1.00 (4 / 4)
```

```
+  538    3.2  0.0  0.0  UnnamedSequence    158   219    (2) + DNAREP1_DM RC/Helitron    533    594    (0)   1
```

```
ANNOTATION EVIDENCE:
  538   3.23 0.00 0.00  UnnamedSequence    158   219      2 + DNAREP1_DM RC/Helitron    533    594      0    
538 3.23 0.00 0.00 UnnamedSequence 158 219 (2) DNAREP1_DM#RC/Helitron 533 594 (0) m_b1s001i1

  UnnamedSequen        158 CACTAGCTGAGTAACGGGTATCAGATAGTCGGGGAACTCGACTATAGCGT 207
                                                 v                         i 
  DNAREP1_DM#RC        533 CACTAGCTGAGTAACGGGTATCTGATAGTCGGGGAACTCGACTATAGCAT 582

  UnnamedSequen        208 TCTCTCTTGTTT 219
                                       
  DNAREP1_DM#RC        583 TCTCTCTTGTTT 594

Matrix = 20p43g.matrix
Kimura (with divCpGMod) = 3.30
Transitions / transversions = 1.00 (1/1)
Gap_init rate = 0.00 (0 / 61), avg. gap size = 0.0 (0 / 0)
```
