## Supplementary material for "Odorant receptor copy number change, co-expression, and positive selection establish peripheral coding differences between fly species": sup. file 3: RM2sequpload_1628782074.out.html

[ close all hsps ]
[ open all hsps ]

```
                                          position in query-                           -position in repeat-
          %    %    %    query                               C matching  repeat        (left)  end   begin  linkage
+ score  div. del. ins.  sequence         begin  end  (left) + repeat    class/family  begin   end   (left) id/graphic
```

```
+  236   11.8  0.0  0.0  UnnamedSequence    146   179  (815) + LINEJ1_DM LINE/I-Jockey   4469   4502  (517)   1
```

```
ANNOTATION EVIDENCE:
  236  11.76 0.00 0.00  UnnamedSequence    146   179    815 + LINEJ1_DM LINE/I-Jockey   4469   4502    517    
236 11.76 0.00 0.00 UnnamedSequence 146 179 (815) LINEJ1_DM#LINE/I-Jockey 4469 4502 (517) m_b1s001i0

  UnnamedSequen        146 TGTTCTACGGCCTTCAGGTATACAGTATTTCTGC 179
                                        v         i i   v    
  LINEJ1_DM#LIN       4469 TGTTCTACGGCCTGCAGGTATACGGCATTGCTGC 4502

Matrix = 20p43g.matrix
Kimura (with divCpGMod) = 9.72
Transitions / transversions = 1.00 (2/2)
Gap_init rate = 0.00 (0 / 33), avg. gap size = 0.0 (0 / 0)
```

```
+   17   18.8  4.0  2.0  UnnamedSequence    210   259  (735) + (AAT)n    Simple_repeat      1     51    (0)   2
```

```
ANNOTATION EVIDENCE:
   17  18.79 4.00 1.96  UnnamedSequence    210   259    735 + (AAT)n    Simple_repeat      1     51      0    
17 18.79 4.00 1.96 UnnamedSequence 210 259 (735) (AAT)n#Simple_repeat 1 51 (0) m_b1s252i0

  UnnamedSequen        210 AATAATAGAAATTTTTA-AATAATAATAATAATACTAGTAAATAGT-ATA 257
                                  iv   vv v -                v  i -    i -   
  (AAT)n#Simple          1 AATAATAATAATAATAATAATAATAATAATAATAATAAT-AATAATAATA 49

  UnnamedSequen        258 AT 259
                             
  (AAT)n#Simple         50 AT 51

Matrix = Unknown
Transitions / transversions = 0.60 (3/5)
Gap_init rate = 0.06 (3 / 49), avg. gap size = 1.00 (3 / 3)
```
