## Supplementary material for "Odorant receptor copy number change, co-expression, and positive selection establish peripheral coding differences between fly species": sup. file 3: RM2sequpload_1628790273.out.html

[ close all hsps ]
[ open all hsps ]

```
                                          position in query-                            -position in repeat-
          %    %    %    query                               C matching   repeat        (left)  end   begin  linkage
+ score  div. del. ins.  sequence         begin  end  (left) + repeat     class/family  begin   end   (left) id/graphic
```

```
+  226   27.9  1.2  2.5  UnnamedSequence    143   223 (2644) + TART-A     LINE/I-Jockey   7323   7402 (8174)   1
```

```
ANNOTATION EVIDENCE:
  226  27.86 1.23 2.50  UnnamedSequence    143   223   2644 + TART-A     LINE/I-Jockey   7323   7402   8174    
226 27.86 1.23 2.50 UnnamedSequence 143 223 (2644) TART-A#LINE/I-Jockey 7323 7402 (8174) m_b1s001i0

  UnnamedSequen        143 AAGATCATTC-ATCAGCGGAGACTGGAACGCAAAACATCCATGGTGGGGT 191
                              v      -  -- i  v        i         vviiii     i
  TART-A#LINE/I       7323 AAGTTCATTCTAT--GTGGCGACTGGAATGCAAAACATAGGCAATGGGGC 7370

  UnnamedSequen        192 TCTGCGATCACCTGTCACCGAGGTAACTTACT 223
                            v i ?vv i       v  v  i v vi   
  TART-A#LINE/I       7371 TGTACNCGCGCCTGTCAACGTGGCACCGCACT 7402

Matrix = 20p43g.matrix
Kimura (with divCpGMod) = 33.83
Transitions / transversions = 1.00 (11/11)
Gap_init rate = 0.04 (3 / 80), avg. gap size = 1.00 (3 / 3)
```

```
+  256   35.1  1.2  1.2  UnnamedSequence    249   421 (2446) + BS         LINE/I-Jockey   2749   2921 (2205)   2
```

```
ANNOTATION EVIDENCE:
  256  35.09 1.16 1.16  UnnamedSequence    249   421   2446 + BS         LINE/I-Jockey   2749   2921   2205    
256 35.09 1.16 1.16 UnnamedSequence 249 421 (2446) BS#LINE/I-Jockey 2749 2921 (2205) m_b1s001i1

  UnnamedSequen        249 GCCAACATCCTGGCGACTGGAGCACCAACATACTATCAAAGCGCATTAAA 298
                             i  v     v  i     vv v  i   viv    vivi  iiiiv i
  BS#LINE/I-Joc       2749 GCTAAGATCCTCGCAACTGGCTCTCCGACAAGGTATCCGTACGTGCCCAG 2798

  UnnamedSequen        299 TCGAAGACCATCCTGTTTAGACTTTGCAATCTATCGCAACATTCCACACG 348
                           i iv vi  v  v  iv    i  i   i v    iiiii  v   v  v
  BS#LINE/I-Joc       2799 CCATACGCCCTCATGCATAGATTTCGCAGTGTATCATGGTATACCAGACC 2848

  UnnamedSequen        349 ACA-AGCTAAACATCAGAGACAGCTGGGACCTTGAATCGGACCACCTGTC 397
                             v-   -  vi  v v v v         i v  v  v  i   i  i 
  BS#LINE/I-Joc       2849 ACCTAGC-AACTATAACACAAAGCTGGGACTTGGATTCTGATCACTTGCC 2897

  UnnamedSequen        398 TCTCATCACTA-CATTAAAAACAGA 421
                              i    i  -    -i i     
  BS#LINE/I-Joc       2898 TCTTATCATTAGCATT-GAGACAGA 2921

Matrix = 20p43g.matrix
Kimura (with divCpGMod) = 44.17
Transitions / transversions = 1.14 (32/28)
Gap_init rate = 0.02 (4 / 172), avg. gap size = 1.00 (4 / 4)
```

```
+ 1238   30.0  3.9  4.0  UnnamedSequence   1040  1829 (1038) + BS3_DM     LINE/I-Jockey    320    960  (830)   3
```

```
ANNOTATION EVIDENCE:
  644  32.11 5.87 5.11  UnnamedSequence   1040  1738   1129 + Jockey2    LINE/I-Jockey   1760   2463    965    
644 32.11 5.87 5.11 UnnamedSequence 1040 1738 (1129) Jockey2#LINE/I-Jockey 1760 2463 (965) m_b1s001i2

  UnnamedSequen       1040 CCATTCCAGATGTCACCTCCAATAAGACCTATCCGTCTGGAAGAAGTCTC 1089
                                i  i   v   vv  viiiiiv   i  vv   v      --  v
  Jockey2#LINE/       1760 CCATTTCAAATGACACGGCCCGCGGATCCTGTCACTCTCGAAGAA--CTG 1807

  UnnamedSequen       1090 AAACATGATACGTACCCT---------CAAAAGACGAAAGGCGCCAGGTC 1130
                                -- ivi  vi   ---------     ----   ii v? i    
  Jockey2#LINE/       1808 AAACA--ACTTGTTTCCTTGTTGAANTCAAAA----AAAAACCNCGGGTC 1851

  UnnamedSequen       1131 ACGACCTAATAAGCAACGCAGTACTTAAAATCCTTCCCAAGAGAGCACTA 1180
                            i     vv vii    iv iiiv v    i   v  vi vvi   v  i
  Jockey2#LINE/       1852 ATGACCTTCTCGACAACAGAACGATAAAAACCCTACCAGACCAAGCTCTG 1901

  UnnamedSequen       1181 TTACTTATCACATTGATATTCAAT-GCGAT--ACTT-AGAGTGCAATACT 1226
                           iv i  v vii i v  v  i  i- ? v --    -  i  ----  - 
  Jockey2#LINE/       1902 CGATTTCTGGTACTCATTTTTAACAGNGTTTTACTTGAGGGT----TA-T 1946

  UnnamedSequen       1227 TTCCCAAAAAT--GGAAGAGTGCTAGAATCAGTATGATTCTAAAGCCAGG 1274
                                    i --      v   v iv  v viv    v  i  i     
  Jockey2#LINE/       1947 TTCCCAAAAGTCTGGAAGACTGCAAACATAATCCTGATACTGAAACCAGG 1996

  UnnamedSequen       1275 AAGCCGG--AACAG-GATCCAAGCTCCTACCGGCCTATCAGTCTCCTGCC 1321
                           i ivv  --     - i -- ii   v  iv i  v     i     v  
  Jockey2#LINE/       1997 GAAAAGGCCAACAGAGGT--AGACTCGTATAGACCAATCAGCCTCCTCCC 2044

  UnnamedSequen       1322 CTCCTTATCG-AAGGTAATGGAAAGGCTGATAGCTTCCCGACTTATAATA 1370
                           i   i - v -  ii v         v i  ----  i i  -- vi  i
  Jockey2#LINE/       2045 TTCCCT-TGGTAAAATTATGGAAAGGATAAT----TCTCAAC--AGGATG 2087

  UnnamedSequen       1371 CATCTAGAAGACAATGA-----TACTAT--------CCCAATGCACCAAT 1407
                            i ------   iv   -----  v   --------  ii  vi-     
  Jockey2#LINE/       2088 CGT------GACGTTGAGCCTGTAGTATTGGCGATACCTGATTT-CCAAT 2130

  UnnamedSequen       1408 TCGGATTCAGAGCTGGCCACAGTACGATTGAGCAACTGCACCGTGTAGTC 1457
                                   v ii  viv  ii v  i?ii        v  iv v  v   
  Jockey2#LINE/       2131 TCGGATTCCGGACTCAACATGGAACANCCGAGCAACTCCATAGAGTTGTC 2180

  UnnamedSequen       1458 AATCATATCCT-GAAGGCCTATGACCATAAAGAATACTGCAACGG-AATC 1505
                              iv iiv  -    v    -  vviv     i  -   iiii - iii
  Jockey2#LINE/       2181 AATTTTGCACTAGAAGCCCTA-GAAAGGAAAGAGTA-TGCGGTAGCAGCT 2228

  UnnamedSequen       1506 TTCCTCGATATTCAACAGGCATTTGATAGAGTTTGG-ATCCCGGGACTAC 1554
                             i  v  i        i  v     ii    v   -    i-     v 
  Jockey2#LINE/       2229 TTTCTGGACATTCAACAAGCTTTTGACGGAGTGTGGCATCCT-GGACTTC 2277

  UnnamedSequen       1555 TAGCCAAAGTCAAAACAGCGCTGCCAGCCAACCTGTTTGAGCTCATCA-G 1603
                             vvi    iv  i v iiv  iv  v  v v  i  iv    ii v - 
  Jockey2#LINE/       2278 TACATAAAGCAAAGAAAATTCTAACACCCCAGCTATTCCAGCTTGTGACG 2327

  UnnamedSequen       1604 ATCGTTTCTCAAACAGAGAGTTT-TGCGT-CCAGTCCAGAGACGCAA--C 1649
                            v v   iv v iv vi? v   -  i  -v iii v vi  i   v-- 
  Jockey2#LINE/       2328 AGCTTTTTGCTAGGACGNACTTTCTGTGTGACGACCGATGGATGCACTTC 2377

  UnnamedSequen       1650 CTCTG-TAAATGTTGCCATTGCAGCTGGCGTACCTCAAGGAAGCGTCTTA 1698
                           v   i-    ----     i v      v              i  v   
  Jockey2#LINE/       2378 GTCTATTAAA----GCCATCGAAGCTGGAGTACCTCAAGGAAGTGTGTTA 2423

  UnnamedSequen       1699 GGACCGATACTTTACTCCATCTATACAGCAGACATCCCCA 1738
                             v  v iv  v          viv v  i     v    
  Jockey2#LINE/       2424 GGCCCTACTCTATACTCCATCTTCTCCGCGGACATGCCCA 2463

Matrix = 20p43g.matrix
Kimura (with divCpGMod) = 39.16
Transitions / transversions = 1.11 (112/101)
Gap_init rate = 0.08 (54 / 698), avg. gap size = 1.43 (77 / 54)

 1238  29.57 3.42 3.74  UnnamedSequence   1187  1829   1038 + BS3_DM     LINE/I-Jockey    320    960    830    
1238 29.57 3.42 3.74 UnnamedSequence 1187 1829 (1038) BS3_DM#LINE/I-Jockey 320 960 (830) m_b1s001i3

  UnnamedSequen       1187 ATCACATTGATATTCAATGCGATACTTAGAGTGCAATACTTTCC-CAAAA 1235
                             i  i  vv       i  v  iv v  iv          i  - v v 
  BS3_DM#LINE/I        320 ATTACGTTTTTATTCAACGCCATGGTGAGGCTGCAATACTTCCCTCCACA 369

  UnnamedSequen       1236 ATGGAAGAGTGCTAGAATCAGTATGATTC-TAAAGCCAGGAAGCCGGAAC 1284
                           i      vvi v  vv  ivvi       -i   v ii i     v   v
  BS3_DM#LINE/I        370 GTGGAAGCTCGGTATTATTTCCATGATTCACAAACCTGGAAAGCCTGAAA 419

  UnnamedSequen       1285 AGGATCCAAGCTCCTACCGGCCTATCAGTCTCCTGCCCTCCTTATCGAAG 1334
                             i i  vi v           v           v  i  vv v      
  BS3_DM#LINE/I        420 AGAACCCTGGGTCCTACCGGCCAATCAGTCTCCTCCCTTCGATCTCGAAG 469

  UnnamedSequen       1335 GTAATGGAAAGGCTGATAGCTTCCCGACTTATAATACATCTAGAAGACAA 1384
                             iv ?  i  i     v   v    iv vi v vivv v i    vv  
  BS3_DM#LINE/I        470 GTGTTNGAGAGACTGATTGCTGCCCGGATGGTCAGGATTATGGAAGCGAA 519

  UnnamedSequen       1385 TGATACTAT-CCCAATGCACCAATTCGGATTCAGAGCTGGCCACAGTACG 1433
                           v i  iiv -   i -   i  i  i  v  iv v     ?   v     
  BS3_DM#LINE/I        520 GGGTATCCTGCCCGA-GCATCAGTTTGGTTTTCGTGCTGGNCACTGTACG 568

  UnnamedSequen       1434 ATTGAGCAACTGCACCGTGTAGTCAATCATATCCTGAAGGCCTATGACCA 1483
                           i v  i     i   v v  i   i v  v     ivv   i vi  vv 
  BS3_DM#LINE/I        569 GTAGAACAACTACACAGAGTGGTCGAGCAAATCCTATCGGCTTTCGAAAA 618

  UnnamedSequen       1484 TAAAGAATACTGCAACGGAATCTTCCTCGATATTCAACAGGCATTTGATA 1533
                           i  i  i          v v       v   i v ivv    i  i   v
  BS3_DM#LINE/I        619 CAAGGAGTACTGCAACGCACTCTTCCTGGATGTACGTGAGGCGTTCGATC 668

  UnnamedSequen       1534 GAGTTTGGATCCCGGGACTACTAGCCAAAGTCAAAACAGCGCTGCCAGCC 1583
                            i  v   vv i v  v? v  ivi    i i  i vvi       v  -
  BS3_DM#LINE/I        669 GGGTGTGGCACTCCGGTNTCCTGCTCAAAATTAAGAATACGCTGCCTGC- 717

  UnnamedSequen       1584 AACC-TG-TTTGAGCTCATCAGATCGTTTCTCAAACAGAG-AGTTTTGCG 1630
                           -   - i-  i iv   v v  i    v    i  v    -  v      
  BS3_DM#LINE/I        718 -ACCATACTTCGGCCTCCTGAGGTCGTATCTCGAAAAGAGAAGATTTGCG 766

  UnnamedSequen       1631 T--CCAGTCCAGAGACGCAACCTCTGT-AAATG---TTGCCATTGCAGCT 1674
                           v-- v v   ----  v ii  -    -     ---v ivii v     i
  BS3_DM#LINE/I        767 GTACGATTCC----ACTCGGCC-CTGTCAAATGAGCATAATGTGGCAGCC 811

  UnnamedSequen       1675 GGCGTACCTCAAGGAAGCGTCTTAGGACCGATACTTTACTCCATCTATAC 1724
                             v     v  i     i  vi v  ?   v i       v --  i   
  BS3_DM#LINE/I        812 GGAGTACCACAGGGAAGTGTACTTGGNCCGCTGCTTTACTGC--CTGTAC 859

  UnnamedSequen       1725 AGC--AGACATCCCCAAGCCAAGCTATTATGAAATGACCTA--TGACAAC 1770
                              --v     v  ?vi    ii -i   -------i    --i i-i  
  BS3_DM#LINE/I        860 AGCTACGACATGCCNCGGCCAGAC-GTTA-------GCCTACCCGG-GAC 900

  UnnamedSequen       1771 AGCAAAATGCTGCTGGCCAGCTTCGCTGACGACGTCTGTTTT-TCT-CAG 1818
                           iv   -----  i      vv  i           v   v i-i  -   
  BS3_DM#LINE/I        901 GTCAA-----TGTTGGCCACATTTGCTGACGACGTGTGTGTCACCTACAG 945

  UnnamedSequen       1819 CTCCT----CGAACA 1829
                           v    ----      
  BS3_DM#LINE/I        946 GTCCTGCTGCGAACA 960

Matrix = 20p43g.matrix
Kimura (with divCpGMod) = 34.05
Transitions / transversions = 0.87 (85/98)
Gap_init rate = 0.06 (38 / 642), avg. gap size = 1.21 (46 / 38)
```

```
+  362   33.4  2.8  4.0  UnnamedSequence   1994  2402  (465) + BS3_DM     LINE/I-Jockey    891   1525  (265)   4
```

```
ANNOTATION EVIDENCE:
  297  35.67 2.05 5.28  UnnamedSequence   1994  2286    581 + HELENA_RT  LINE/I-Jockey    891   1174    143    
297 35.67 2.05 5.28 UnnamedSequence 1994 2286 (581) HELENA_RT#LINE/I-Jockey 891 1174 (143) m_b1s001i4

  UnnamedSequen       1994 GCAAGCAACAAAGGCGAAATACCTAGGCCTCACACTGGACAAACGCCTAA 2043
                                     ---------  i     iv    v     i     v  v 
  HELENA_RT#LIN        891 GCAAGCAACA---------TATCTAGGTATCACCCTGGATAAACGGCTCA 931

  UnnamedSequen       2044 CCTTCCGAGATCATATTGCCAGAGT-C-GTCAA--AATGTGCAACCTAAA 2089
                               i--i ii ii  iv v i ii- -  v  --      -iv vi i 
  HELENA_RT#LIN        932 CCTTT--GGGCCGCATCTCAAAAACACAGTAAAGAAATGTG-GTCACAGA 978

  UnnamedSequen       2090 GCGTAACCAACTATTCTGGATGCTAAATAAGAAAAGCAAACTACCTTTAA 2139
                           v ii   - i  ivvv   v vv i    ii ii    vv  vi vi i 
  HELENA_RT#LIN        979 TCACAAC-AGCTGAGATGGCTCATGAATAGAAGGAGCACTCTTTCGCTGA 1027

  UnnamedSequen       2140 GATGCAAGCGTCAG--ATTTATCAGCAAATTATCGCACCTACTTGGAGAT 2187
                            i     iv --  --i v   vv   vvv     i   v iv   vv  
  HELENA_RT#LIN       1028 GGTGCAAAAG--AGCTGTGTATGCGCACTGTATCGTACCGATGTGGTTAT 1075

  UnnamedSequen       2188 ACGGCATCCAAATTTGGGGAGTGGCGGCTGCGTCCCACCGTAAACGATTC 2237
                               v     i         i v  i  i vi  iv iii      vv  
  HELENA_RT#LIN       1076 ACGGGATCCAGATTTGGGGAATTGCAGCCGAATCTAATTATAAACGTATC 1125

  UnnamedSequen       2238 CAAACCGTCCAGAACAAAACATTAAGACAAATCACTGGCTGTGACTGGT 2286
                             iiivi v  i  ivivi  i  v       v  iii    vv     
  HELENA_RT#LIN       1126 CAGGTGATGCAAAATCGCGCACTACGACAAATAACCAACTGTCCCTGGT 1174

Matrix = 20p43g.matrix
Kimura (with divCpGMod) = 46.94
Transitions / transversions = 1.20 (54/45)
Gap_init rate = 0.07 (19 / 292), avg. gap size = 1.11 (21 / 19)

  362  31.95 3.21 3.21  UnnamedSequence   2154  2402    465 + BS3_DM     LINE/I-Jockey   1277   1525    265    
362 31.95 3.21 3.21 UnnamedSequence 2154 2402 (465) BS3_DM#LINE/I-Jockey 1277 1525 (265) m_b1s001i5

  UnnamedSequen       2154 ATTTATCAGCAAATTATCGCACCTACTTGGAGATACGGCATCCAAATTTG 2203
                                iv          v  i  v iv     i  i  ivvi  i     
  BS3_DM#LINE/I       1277 ATTTACAAGCAAATTATAGCGCCAATATGGAGGTATGGTTGTCAGATTTG 1326

  UnnamedSequen       2204 GGGAGTGGCGGCTGCGTC--CCACCGTAAACGATTCCAAACCGTCCAGAA 2251
                              vv    --i    v --   vvv viv  vv i  ii i i   i  
  BS3_DM#LINE/I       1327 GGGCTTGGC--TTGCGACAGCCAAATTCGCCGCATTCAGGCTGCCCAAAA 1374

  UnnamedSequen       2252 CAAAA-CATTAAGACAAATCACTGGCTGTGACTGGTTCGTTAGCGGCCAA 2300
                                - iii i   -i  i  i     i  v    v   v  vii vv 
  BS3_DM#LINE/I       1375 CAAAATCGCCAGGAC-GATTACCGGCTGCGAATGGTACGTAAGGAACACA 1423

  UnnamedSequen       2301 ACGCTTCACAATGATTTAAACCTGAGCC-TTG--TTGAAGACCA-AATCT 2346
                             v  v     v  ii v  v  -    -i  --  vi  i v -  v v
  BS3_DM#LINE/I       1424 ACCCTGCACAAAGACCTCAAGCT-AGCCACTGTCTTCGAGGCAATAAACA 1472

  UnnamedSequen       2347 CGTTCTTCTCAAGCAGATACAACGACCGACTAACTGC-CCACTGTAATCG 2395
                           i iv  i ---   v i   v     v i   iv-  -    v v     
  BS3_DM#LINE/I       1473 TGCACTCC---AGCCGGTACCACGACAGGCTAGA-GCGCCACAGAAATCG 1518

  UnnamedSequen       2396 TCTCGCC 2402
                           i  v   
  BS3_DM#LINE/I       1519 CCTAGCC 1525

Matrix = 20p43g.matrix
Kimura (with divCpGMod) = 37.44
Transitions / transversions = 0.97 (38/39)
Gap_init rate = 0.06 (14 / 248), avg. gap size = 1.14 (16 / 14)
```

```
+  960    9.5  0.5  3.8  UnnamedSequence   2666  2859    (8) + DNAREP1_DM RC/Helitron      263    509   (85)   5
```

```
ANNOTATION EVIDENCE:
  861  12.61 0.00 5.66  UnnamedSequence   2666  2833     34 + DNAREP1_DM RC/Helitron      351    509     85    
861 12.61 0.00 5.66 UnnamedSequence 2666 2833 (34) DNAREP1_DM#RC/Helitron 351 509 (85) m_b1s001i6

  UnnamedSequen       2666 ATTAGTCTTGTAATTTTCTATCGATTTACCAAAAAACTTTTTGCCACGTC 2715
                                        v             i                    i 
  DNAREP1_DM#RC        351 ATTAGTCTTGTAAATTTCTATCGATTTGCCAAAAAACTTTTTGCCACGCC 400

  UnnamedSequen       2716 CACTCTAACGTCTACAAACCGCCCAAAACTGCCACGCCCACACTTTTGAA 2765
                                     i ivv  i                                
  DNAREP1_DM#RC        401 CACTCTAACGCCCTAAAGCCGCCCAAAACTGCCACGCCCACACTTTTGAA 450

  UnnamedSequen       2766 AAATGTTTTGAAATTTTTTTCATTTTTGTATTATTCTTGTAATTTTCTAT 2815
                           ?   -    i  v     i     ------      ii-i  - v     
  DNAREP1_DM#RC        451 NAAT-TTTTAAATTTTTTCTCATT------TTATTCCC-CAA-TATCTAT 491

  UnnamedSequen       2816 CGATTTGTCAAAAAACTT 2833
                               v vi  i    v  
  DNAREP1_DM#RC        492 CGATATCCCAGAAAAATT 509

Matrix = 20p43g.matrix
Kimura (with divCpGMod) = 13.98
Transitions / transversions = 1.50 (12/8)
Gap_init rate = 0.05 (9 / 167), avg. gap size = 1.00 (9 / 9)

  960   8.73 0.65 3.33  UnnamedSequence   2706  2859      8 + DNAREP1_DM RC/Helitron      263    412    182    
960 8.73 0.65 3.33 UnnamedSequence 2706 2859 (8) DNAREP1_DM#RC/Helitron 263 412 (182) m_b1s001i7

  UnnamedSequen       2706 TTGCCACGTCCACTCTAACGTCTACAAACCGCCCAAAACTGCCACGCCCA 2755
                                   i           i i                           
  DNAREP1_DM#RC        263 TTGCCACGCCCACTCTAACGCCCACAAACCGCCCAAAACTGCCACGCCCA 312

  UnnamedSequen       2756 CACTTTTGAAAAA-TGTTTTGAAATTTTTTTCATTTTTGTATTATTCTTG 2804
                                        - v     --  v        v?  v     v     
  DNAREP1_DM#RC        313 CACTTTTGAAAAANTTTTTTG--ATATTTTTCATANTTTTATTAGTCTTG 360

  UnnamedSequen       2805 TAATTTTCTATCGATTTGTCAAAAAACTTTCTGCCACGCCCATATATATA 2854
                              v              i           i           i--- v  
  DNAREP1_DM#RC        361 TAAATTTCTATCGATTTGCCAAAAAACTTTTTGCCACGCCCAC---TCTA 407

  UnnamedSequen       2855 ACGCC 2859
                                
  DNAREP1_DM#RC        408 ACGCC 412

Matrix = 20p43g.matrix
Kimura (with divCpGMod) = 9.35
Transitions / transversions = 0.86 (6/7)
Gap_init rate = 0.04 (6 / 153), avg. gap size = 1.00 (6 / 6)
```
