## Supplementary material for "Odorant receptor copy number change, co-expression, and positive selection establish peripheral coding differences between fly species": sup. file 3: RM2sequpload_1628858592.out.html

[ close all hsps ]
[ open all hsps ]

```
                                          position in query-                             -position in repeat-
          %    %    %    query                               C matching    repeat        (left)  end   begin  linkage
+ score  div. del. ins.  sequence         begin  end  (left) + repeat      class/family  begin   end   (left) id/graphic
```

```
+ 3295    7.9  5.5  1.8  UnnamedSequence      2   136  (686) + DNAREP1_DM  RC/Helitron        1    149  (445)   1
```

```
ANNOTATION EVIDENCE:
 3295   7.93 5.53 1.78  UnnamedSequence      2   136    686 + DNAREP1_DM  RC/Helitron        1    149    445    
3295 7.93 5.53 1.78 UnnamedSequence 2 136 (686) DNAREP1_DM#RC/Helitron 1 149 (445) m_b1s001i0

  UnnamedSequen          2 TTATACCCGTTACTCGTAGAGTAACAGGGTATACGAGATTCGTTGAAAAG 51
                                                   v         v               
  DNAREP1_DM#RC          1 TTATACCCGTTACTCGTAGAGTAAAAGGGTATACTAGATTCGTTGAAAAG 50

  UnnamedSequen         52 TATGTAACAGGCAGAAGGAAGCGTTTCCGACCATATCAAGCATAT----- 96
                                                               v   i    -----
  DNAREP1_DM#RC         51 TATGTAACAGGCAGAAGGAAGCGTTTCCGACCATATAAAGTATATNCGCG 100

  UnnamedSequen         97 ---------ATATTCTTGATCAGGATCAATAGCCGAGTCGATCTGGCCA 136
                           ---------                                        
  DNAREP1_DM#RC        101 CGCGCGCGTATATTCTTGATCAGGATCAATAGCCGAGTCGATCTGGCCA 149

Matrix = 20p43g.matrix
Kimura (with divCpGMod) = 8.27
Transitions / transversions = 0.33 (1/3)
Gap_init rate = 0.01 (2 / 134), avg. gap size = 7.00 (14 / 2)
```

```
+   33    5.2  0.0  0.0  UnnamedSequence    137   176  (646) + (TGTCCGTC)n Simple_repeat      1     40    (0)   2
```

```
ANNOTATION EVIDENCE:
   33   5.18 0.00 0.00  UnnamedSequence    137   176    646 + (TGTCCGTC)n Simple_repeat      1     40      0    
33 5.18 0.00 0.00 UnnamedSequence 137 176 (646) (TGTCCGTC)n#Simple_repeat 1 40 (0) c_b1s251i0

  UnnamedSequen        137 TGTCCGTCTGTCCGTCCGTCCGTCTGTCCTTCTGTCCGTC 176
                                           i            v          
  (TGTCCGTC)n#S          1 TGTCCGTCTGTCCGTCTGTCCGTCTGTCCGTCTGTCCGTC 40

Matrix = Unknown
Transitions / transversions = 1.00 (1/1)
Gap_init rate = 0.00 (0 / 39), avg. gap size = 0.0 (0 / 0)
```

```
+ 3295    9.5  3.9  2.0  UnnamedSequence    177   701  (121) + DNAREP1_DM  RC/Helitron      150    506   (88)   1
```

```
ANNOTATION EVIDENCE:
 3295   7.93 5.53 1.78  UnnamedSequence    177   528    294 + DNAREP1_DM  RC/Helitron      150    505     89    
3295 7.93 5.53 1.78 UnnamedSequence 177 528 (294) DNAREP1_DM#RC/Helitron 150 505 (89) m_b1s001i0

  UnnamedSequen        177 -----------CGTATGAATGTCGAGATCTCAGGAACTACAAAAGCTAGA 215
                           -----------        i                   i          
  DNAREP1_DM#RC        150 GTCCGTCTGTCCGTATGAACGTCGAGATCTCAGGAACTATAAAAGCTAGA 199

  UnnamedSequen        216 AAGTTAAGATTAAGCATACAAACTCCAGAGACATAGAAGCAGCGCAAGTT 265
                            i   i     ?        i i  i           v            
  DNAREP1_DM#RC        200 AGGTTGAGATTNAGCATACAGATTCTAGAGACATAGACGCAGCGCAAGTT 249

  UnnamedSequen        266 TGTCGATTCATGTTGCCACGCCCATTCTAACGCCCACAAACCGCCCAAAA 315
                              i  ii                i                         
  DNAREP1_DM#RC        250 TGTTGACCCATGTTGCCACGCCCACTCTAACGCCCACAAACCGCCCAAAA 299

  UnnamedSequen        316 CTGCCACGCCCACACTTTTGAAAAA-TGTTTTGAAATTTTTTCATTTTTG 364
                                                    - v      v -        v?  v
  DNAREP1_DM#RC        300 CTGCCACGCCCACACTTTTGAAAAANTTTTTTGATA-TTTTTCATANTTT 348

  UnnamedSequen        365 TATTAGTCTTGTAATTTTCTATCGATTTACCAAAAAACTTTTTGCCACGC 414
                                         v             i                     
  DNAREP1_DM#RC        349 TATTAGTCTTGTAAATTTCTATCGATTTGCCAAAAAACTTTTTGCCACGC 398

  UnnamedSequen        415 CCACTCTAACGCCCTCAAACCGCCCAAAGCTGATACGCCCACACTTTTGA 464
                                          v  i         i   vi                
  DNAREP1_DM#RC        399 CCACTCTAACGCCCTAAAGCCGCCCAAAACTGCCACGCCCACACTTTTGA 448

  UnnamedSequen        465 AAAATGTTTTGATTTTTT-TCATTTTTATATTGGTCTTGTGAATTTCTAT 513
                            ?   v   vi       -      ---    vvi -----   v     
  DNAREP1_DM#RC        449 ANAATTTTTAAATTTTTTCTCATTT---TATTCCCC-----AATATCTAT 490

  UnnamedSequen        514 CGATTTGCCAAGAAA 528
                               v v   ii   
  DNAREP1_DM#RC        491 CGATATCCCAGAAAA 505

Matrix = 20p43g.matrix
Kimura (with divCpGMod) = 8.27
Transitions / transversions = 1.27 (19/15)
Gap_init rate = 0.03 (12 / 351), avg. gap size = 1.83 (22 / 12)

 1430  12.77 0.40 3.28  UnnamedSequence    406   656    166 + DNAREP1_DM  RC/Helitron      263    506     88    
1430 12.77 0.40 3.28 UnnamedSequence 406 656 (166) DNAREP1_DM#RC/Helitron 263 506 (88) m_b1s001i1

  UnnamedSequen        406 TTGCCACGCCCACTCTAACGCCCTCAAACCGCCCAAAGCTGATACGCCCA 455
                                                  v             i   vi       
  DNAREP1_DM#RC        263 TTGCCACGCCCACTCTAACGCCCACAAACCGCCCAAAACTGCCACGCCCA 312

  UnnamedSequen        456 CACTTTTGAAAAA-TGTTTTGATTTTTTTCATTTTTATATTGGTCTTGTG 504
                                        - v       v        v?  v    i       i
  DNAREP1_DM#RC        313 CACTTTTGAAAAANTTTTTTGATATTTTTCATANTTTTATTAGTCTTGTA 362

  UnnamedSequen        505 AATTTCTATCGATTTGCCAAGAAACGTTCTGCCACGCCCACTCTAACGCC 554
                                               i    v  i                     
  DNAREP1_DM#RC        363 AATTTCTATCGATTTGCCAAAAAACTTTTTGCCACGCCCACTCTAACGCC 412

  UnnamedSequen        555 CTTAAACCGCCCAAAGCTGATACGCCCACACTTTTGAAAAATGTTTTGAT 604
                             v  i         i   vi                 ?   v   vi  
  DNAREP1_DM#RC        413 CTAAAGCCGCCCAAAACTGCCACGCCCACACTTTTGAANAATTTTTAAAT 462

  UnnamedSequen        605 ATTTTTTCATTTTTATATTGGTCTTGTGAATTTCTATCGATTTGCCAAGA 654
                           v    i      ---    vvi -----   v         v v   ii 
  DNAREP1_DM#RC        463 TTTTTCTCATTT---TATTCCCC-----AATATCTATCGATATCCCAGAA 504

  UnnamedSequen        655 AA 656
                             
  DNAREP1_DM#RC        505 AA 506

Matrix = 20p43g.matrix
Kimura (with divCpGMod) = 14.14
Transitions / transversions = 0.82 (14/17)
Gap_init rate = 0.04 (9 / 250), avg. gap size = 1.00 (9 / 9)

 1052  11.97 0.60 0.60  UnnamedSequence    534   701    121 + DNAREP1_DM  RC/Helitron      264    431    163    
1052 11.97 0.60 0.60 UnnamedSequence 534 701 (121) DNAREP1_DM#RC/Helitron 264 431 (163) m_b1s001i2

  UnnamedSequen        534 TGCCACGCCCACTCTAACGCCCTTAAACCGCCCAAAGCTGATACGCCCAC 583
                                                 vi            i   vi        
  DNAREP1_DM#RC        264 TGCCACGCCCACTCTAACGCCCACAAACCGCCCAAAACTGCCACGCCCAC 313

  UnnamedSequen        584 ACTTTTGAAAAA-TGTTTTGATATTTTTTCATTTTTATATTGGTCTTGTG 632
                                       - v        -        v?  v    i       i
  DNAREP1_DM#RC        314 ACTTTTGAAAAANTTTTTTGATA-TTTTTCATANTTTTATTAGTCTTGTA 362

  UnnamedSequen        633 AATTTCTATCGATTTGCCAAGAAACGTTCTGCCACGACCACTATAACGCC 682
                                               i    v  i       v     v       
  DNAREP1_DM#RC        363 AATTTCTATCGATTTGCCAAAAAACTTTTTGCCACGCCCACTCTAACGCC 412

  UnnamedSequen        683 TACAAACCGCCAAAAACTG 701
                           ivv  i     v       
  DNAREP1_DM#RC        413 CTAAAGCCGCCCAAAACTG 431

Matrix = 20p43g.matrix
Kimura (with divCpGMod) = 13.15
Transitions / transversions = 0.82 (9/11)
Gap_init rate = 0.01 (2 / 167), avg. gap size = 1.00 (2 / 2)
```

```
+  538    3.2  0.0  0.0  UnnamedSequence    727   788   (34) + DNAREP1_DM  RC/Helitron      533    594    (0)   1
```

```
ANNOTATION EVIDENCE:
  538   3.23 0.00 0.00  UnnamedSequence    727   788     34 + DNAREP1_DM  RC/Helitron      533    594      0    
538 3.23 0.00 0.00 UnnamedSequence 727 788 (34) DNAREP1_DM#RC/Helitron 533 594 (0) m_b1s001i3

  UnnamedSequen        727 CACTAGCTGAGTAACGGGTATCAGATAGTCGGGGAACTCGACTATAGCGT 776
                                                 v                         i 
  DNAREP1_DM#RC        533 CACTAGCTGAGTAACGGGTATCTGATAGTCGGGGAACTCGACTATAGCAT 582

  UnnamedSequen        777 TCTCTCTTGTTT 788
                                       
  DNAREP1_DM#RC        583 TCTCTCTTGTTT 594

Matrix = 20p43g.matrix
Kimura (with divCpGMod) = 3.30
Transitions / transversions = 1.00 (1/1)
Gap_init rate = 0.00 (0 / 61), avg. gap size = 0.0 (0 / 0)
```
