## Supplementary material for "Odorant receptor copy number change, co-expression, and positive selection establish peripheral coding differences between fly species": sup. file 3: RM2sequpload_1628789679.out.html

[ close all hsps ]
[ open all hsps ]

```
                                          position in query-                          -position in repeat-
          %    %    %    query                               C matching   repeat      (left)  end   begin  linkage
+ score  div. del. ins.  sequence         begin  end  (left) + repeat     class/famil begin   end   (left) id/graphic
```

```
+ 3269    9.3  2.3  4.7  UnnamedSequence      1   818   (86) + DNAREP1_DM RC/Helitron      1    507   (87)   1
```

```
ANNOTATION EVIDENCE:
 3269   8.00 3.31 5.15  UnnamedSequence      1   514    390 + DNAREP1_DM RC/Helitron      1    505     89    
3269 8.00 3.31 5.15 UnnamedSequence 1 514 (390) DNAREP1_DM#RC/Helitron 1 505 (89) m_b1s001i0

  UnnamedSequen          1 TTATACCCGTTACTCGTAGAGTAAAAGGGTATACTAGATTCGTTGAAAAG 50
                                                                             
  DNAREP1_DM#RC          1 TTATACCCGTTACTCGTAGAGTAAAAGGGTATACTAGATTCGTTGAAAAG 50

  UnnamedSequen         51 TATGTAACAGGCAGAAGAAAGCGTTTCCGACCATATAAAGTATAT----- 95
                                            i                           -----
  DNAREP1_DM#RC         51 TATGTAACAGGCAGAAGGAAGCGTTTCCGACCATATAAAGTATATNCGCG 100

  UnnamedSequen         96 ---------ATAATCTTGATCAGGATCAATAGCCG-GTCGATCTGGCCAT 135
                           ---------   v                      -              
  DNAREP1_DM#RC        101 CGCGCGCGTATATTCTTGATCAGGATCAATAGCCGAGTCGATCTGGCCAT 150

  UnnamedSequen        136 GTCCGTCTGTCCGTCCGTCTGTCCGGCCGTATGAACGTCGAGATCTCAGG 185
                                         ----------------                    
  DNAREP1_DM#RC        151 GTCCGTCTGTCCGT----------------ATGAACGTCGAGATCTCAGG 184

  UnnamedSequen        186 AACTACTAAAGCTAGAAAGTTGAGATTAAGCATACAAACTCCAGGGACAT 235
                                iv          i         ?        i i  i  i     
  DNAREP1_DM#RC        185 AACTATAAAAGCTAGAAGGTTGAGATTNAGCATACAGATTCTAGAGACAT 234

  UnnamedSequen        236 AGAAGCAGGGCAAGTTTGTCGATTCATGTTGCCACGCCCATTCTAACGCC 285
                              v    v          i  ii                i         
  DNAREP1_DM#RC        235 AGACGCAGCGCAAGTTTGTTGACCCATGTTGCCACGCCCACTCTAACGCC 284

  UnnamedSequen        286 CACAAACCGCCCAAAACTGCCACGCCCACACTTTTGAAAAA-TGTTTTGA 334
                                                                    - v      
  DNAREP1_DM#RC        285 CACAAACCGCCCAAAACTGCCACGCCCACACTTTTGAAAAANTTTTTTGA 334

  UnnamedSequen        335 AATTTTTTCATTTTTGTATTAGTCTTGTAATTTTCTATCGATTTACCAAA 384
                           v -        v?  v              v             i     
  DNAREP1_DM#RC        335 TA-TTTTTCATANTTTTATTAGTCTTGTAAATTTCTATCGATTTGCCAAA 383

  UnnamedSequen        385 AAATTTTTTGCCACGCCCACTCTAACGCCCTCAAACCGCCCAAAGCTGCT 434
                              i                           v  i         i    i
  DNAREP1_DM#RC        384 AAACTTTTTGCCACGCCCACTCTAACGCCCTAAAGCCGCCCAAAACTGCC 433

  UnnamedSequen        435 ACGCCCACACTTTTG-ATATTTGTTTTGATATTTTGTCATTTTTATATTA 483
                                          - ? v  -   vi  v    v      ---    v
  DNAREP1_DM#RC        434 ACGCCCACACTTTTGAANAATT-TTTAAATTTTTTCTCATTT---TATTC 479

  UnnamedSequen        484 GTCTTGTAAATTTCTATCGATTGGCCAAAAA 514
                           vi -----   v         vvv   i   
  DNAREP1_DM#RC        480 CCC-----AATATCTATCGATATCCCAGAAA 505

Matrix = 20p43g.matrix
Kimura (with divCpGMod) = 8.52
Transitions / transversions = 0.95 (19/20)
Gap_init rate = 0.06 (31 / 513), avg. gap size = 1.39 (43 / 31)

 1460   9.89 0.79 4.08  UnnamedSequence    392   644    260 + DNAREP1_DM RC/Helitron    263    507     87    
1460 9.89 0.79 4.08 UnnamedSequence 392 644 (260) DNAREP1_DM#RC/Helitron 263 507 (87) m_b1s001i1

  UnnamedSequen        392 TTGCCACGCCCACTCTAACGCCCTCAAACCGCCCAAAGCTGCTACGCCCA 441
                                                  v             i    i       
  DNAREP1_DM#RC        263 TTGCCACGCCCACTCTAACGCCCACAAACCGCCCAAAACTGCCACGCCCA 312

  UnnamedSequen        442 CACTTTTG--ATATTTGTTTTGATATTTTGTCATTTTTATATTAGTCTTG 489
                                   -- v ?  -            -    v?  v           
  DNAREP1_DM#RC        313 CACTTTTGAAAAANTT-TTTTGATATTTT-TCATANTTTTATTAGTCTTG 360

  UnnamedSequen        490 TAAATTTCTATCGATTGGCCAAAAAGCTTTTTGCCACGCCCACTCTAACG 539
                                           v        i                        
  DNAREP1_DM#RC        361 TAAATTTCTATCGATTTGCCAAAAAACTTTTTGCCACGCCCACTCTAACG 410

  UnnamedSequen        540 CCCTCAAACCGCCCAAAGCTGATACGCCCACACTTTTGAAAAATGTTTTG 589
                               v  i         i   vi                 ?   -    i
  DNAREP1_DM#RC        411 CCCTAAAGCCGCCCAAAACTGCCACGCCCACACTTTTGAANAAT-TTTTA 459

  UnnamedSequen        590 ATTTTTTTTTCATTTTATATTGGTCTTGTGAATTTCTATCGATTTGCCAA 639
                            v      i         -----  i--iv   v         v v   i
  DNAREP1_DM#RC        460 AATTTTTTCTCATTTTAT-----TCC--CCAATATCTATCGATATCCCAG 502

  UnnamedSequen        640 AGAAA 644
                            i   
  DNAREP1_DM#RC        503 AAAAA 507

Matrix = 20p43g.matrix
Kimura (with divCpGMod) = 10.76
Transitions / transversions = 1.00 (12/12)
Gap_init rate = 0.04 (11 / 252), avg. gap size = 1.09 (12 / 11)

 1346  12.78 0.39 4.51  UnnamedSequence    520   773    131 + DNAREP1_DM RC/Helitron    263    506     88    
1346 12.78 0.39 4.51 UnnamedSequence 520 773 (131) DNAREP1_DM#RC/Helitron 263 506 (88) m_b1s001i2

  UnnamedSequen        520 TTGCCACGCCCACTCTAACGCCCTCAAACCGCCCAAAGCTGATACGCCCA 569
                                                  v             i   vi       
  DNAREP1_DM#RC        263 TTGCCACGCCCACTCTAACGCCCACAAACCGCCCAAAACTGCCACGCCCA 312

  UnnamedSequen        570 CACTTTTGAAAAA-TGTTTTGATTTTTTTTTCATTTTATATTGGTCTTGT 618
                                        - v       v     ivi ?   -    i       
  DNAREP1_DM#RC        313 CACTTTTGAAAAANTTTTTTGATATTTTTCATANTTT-TATTAGTCTTGT 361

  UnnamedSequen        619 GAATTTCTATCGATTTGCCAAAGAAACGTTCTGCCACGCCCACTCTAACG 668
                           i                     -    v  i                   
  DNAREP1_DM#RC        362 AAATTTCTATCGATTTGCCAAA-AAACTTTTTGCCACGCCCACTCTAACG 410

  UnnamedSequen        669 CCCTTTAAACCGCCCAAAGCTGATACGCCCACACTTTTGAAAAATGTTTT 718
                              - v  i         i   vi                 ?   v   v
  DNAREP1_DM#RC        411 CCC-TAAAGCCGCCCAAAACTGCCACGCCCACACTTTTGAANAATTTTTA 459

  UnnamedSequen        719 GATATTTTTTCATTTTTATATTGGTCTTGTGAATTTCTATCGATTTGCCA 768
                           i  v    i      ---    vvi -----   v         v v   
  DNAREP1_DM#RC        460 AATTTTTTCTCATTT---TATTCCCC-----AATATCTATCGATATCCCA 501

  UnnamedSequen        769 AGAAA 773
                           ii   
  DNAREP1_DM#RC        502 GAAAA 506

Matrix = 20p43g.matrix
Kimura (with divCpGMod) = 14.15
Transitions / transversions = 0.94 (15/16)
Gap_init rate = 0.05 (12 / 253), avg. gap size = 1.00 (12 / 12)

 1022  11.97 0.59 1.19  UnnamedSequence    650   818     86 + DNAREP1_DM RC/Helitron    264    431    163    
1022 11.97 0.59 1.19 UnnamedSequence 650 818 (86) DNAREP1_DM#RC/Helitron 264 431 (163) m_b1s001i3

  UnnamedSequen        650 TGCCACGCCCACTCTAACGCCCTTTAAACCGCCCAAAGCTGATACGCCCA 699
                                                 vi-            i   vi       
  DNAREP1_DM#RC        264 TGCCACGCCCACTCTAACGCCCAC-AAACCGCCCAAAACTGCCACGCCCA 312

  UnnamedSequen        700 CACTTTTGAAAAA-TGTTTTGATATTTTTTCATTTTTATATTGGTCTTGT 748
                                        - v        -        v?  v    i       
  DNAREP1_DM#RC        313 CACTTTTGAAAAANTTTTTTGATA-TTTTTCATANTTTTATTAGTCTTGT 361

  UnnamedSequen        749 GAATTTCTATCGATTTGCCAAGAAACGTTCTGCCACGACCACTATAACGC 798
                           i                    i    v  i       v     v      
  DNAREP1_DM#RC        362 AAATTTCTATCGATTTGCCAAAAAACTTTTTGCCACGCCCACTCTAACGC 411

  UnnamedSequen        799 CTACAAACCGCCAAAAACTG 818
                            ivv  i     v       
  DNAREP1_DM#RC        412 CCTAAAGCCGCCCAAAACTG 431

Matrix = 20p43g.matrix
Kimura (with divCpGMod) = 13.15
Transitions / transversions = 0.82 (9/11)
Gap_init rate = 0.02 (3 / 168), avg. gap size = 1.00 (3 / 3)
```

```
+  529    3.3  0.0  0.0  UnnamedSequence    844   904    (0) + DNAREP1_DM RC/Helitron    533    593    (1)   1
```

```
ANNOTATION EVIDENCE:
  529   3.28 0.00 0.00  UnnamedSequence    844   904      0 + DNAREP1_DM RC/Helitron    533    593      1    
529 3.28 0.00 0.00 UnnamedSequence 844 904 (0) DNAREP1_DM#RC/Helitron 533 593 (1) m_b1s001i4

  UnnamedSequen        844 CACTAGCTGAGTAACGGGTATCAGATAGTCGGGGAACTCGACTATAGCGT 893
                                                 v                         i 
  DNAREP1_DM#RC        533 CACTAGCTGAGTAACGGGTATCTGATAGTCGGGGAACTCGACTATAGCAT 582

  UnnamedSequen        894 TCTCTCTTGTT 904
                                      
  DNAREP1_DM#RC        583 TCTCTCTTGTT 593

Matrix = 20p43g.matrix
Kimura (with divCpGMod) = 3.35
Transitions / transversions = 1.00 (1/1)
Gap_init rate = 0.00 (0 / 60), avg. gap size = 0.0 (0 / 0)
```
