## Supplementary material for "Odorant receptor copy number change, co-expression, and positive selection establish peripheral coding differences between fly species": sup. file 3: RM2sequpload_1628857874.out.html

[ close all hsps ]
[ open all hsps ]

```
                                          position in query-                          -position in repeat-
          %    %    %    query                               C matching   repeat      (left)  end   begin  linkage
+ score  div. del. ins.  sequence         begin  end  (left) + repeat     class/famil begin   end   (left) id/graphic
```

```
+  560    8.3  0.0  0.0  UnnamedSequence    246   317  (988) C DNAREP1_DM RC/Helitron    (0)    594    523   1
```

```
ANNOTATION EVIDENCE:
  560   8.33 0.00 0.00  UnnamedSequence    246   317    988 C DNAREP1_DM RC/Helitron    523    594      0    
560 8.33 0.00 0.00 UnnamedSequence 246 317 (988) C DNAREP1_DM#RC/Helitron (0) 594 523 m_b1s001i0

  UnnamedSequen        246 AAACAAGAGAGAACGCTATAGTCGAGTTCCTCGACTATCTGATACCCGTT 295
                                        i                i        v          
C DNAREP1_DM#RC        594 AAACAAGAGAGAATGCTATAGTCGAGTTCCCCGACTATCAGATACCCGTT 545

  UnnamedSequen        296 ACTCAGCTAGTGAAAGTGCGAA 317
                                       vi i      
C DNAREP1_DM#RC        544 ACTCAGCTAGTGTGAATGCGAA 523

Matrix = 20p43g.matrix
Kimura (with divCpGMod) = 8.91
Transitions / transversions = 2.00 (4/2)
Gap_init rate = 0.00 (0 / 71), avg. gap size = 0.0 (0 / 0)
```

```
+ 2635    8.7  9.4  4.3  UnnamedSequence    333   979  (326) C DNAREP1_DM RC/Helitron   (85)    509      1   1
```

```
ANNOTATION EVIDENCE:
 1100   8.82 1.16 1.16  UnnamedSequence    333   504    801 C DNAREP1_DM RC/Helitron    263    434    160    
1100 8.82 1.16 1.16 UnnamedSequence 333 504 (801) C DNAREP1_DM#RC/Helitron (160) 434 263 m_b1s001i1

  UnnamedSequen        333 TGACAGTTTTTGGCGGTTTGTGAGCGTTAGAGTGGGCGTGGCAAAAAGTT 382
                             i       v     i  vv i                           
C DNAREP1_DM#RC        434 TGGCAGTTTTGGGCGGCTTTAGGGCGTTAGAGTGGGCGTGGCAAAAAGTT 385

  UnnamedSequen        383 TTTTGGCAAATCGATAGAAATTTAGAAGACTAATACAAAAATGAAAAAAT 432
                                                   v          v  ?v        - 
C DNAREP1_DM#RC        384 TTTTGGCAAATCGATAGAAATTTACAAGACTAATAAAANTATGAAAAA-T 336

  UnnamedSequen        433 ATCATAACA-TTTTTCAAA-GCGTGGGCGTGACAGCTTTGGGGCGGTTTT 480
                               v  v -         - i         i   i      -      v
C DNAREP1_DM#RC        335 ATCAAAAAANTTTTTCAAAAGTGTGGGCGTGGCAGTTTTGGG-CGGTTTG 287

  UnnamedSequen        481 TGGGCGTTAGAGTGGGCGTGGCAA 504
                                                   
C DNAREP1_DM#RC        286 TGGGCGTTAGAGTGGGCGTGGCAA 263

Matrix = 20p43g.matrix
Kimura (with divCpGMod) = 9.45
Transitions / transversions = 0.67 (6/9)
Gap_init rate = 0.02 (4 / 171), avg. gap size = 1.00 (4 / 4)

  809  14.61 9.62 4.59  UnnamedSequence    378   585    720 C DNAREP1_DM RC/Helitron    292    509     85    
809 14.61 9.62 4.59 UnnamedSequence 378 585 (720) C DNAREP1_DM#RC/Helitron (85) 509 292 m_b1s001i2

  UnnamedSequen        378 AAGTTTTTTGGCAAATCGATAGAAATTTAGAAGACTAATACAAAAATGAA 427
                             v    i   v v         v   -- ii  v   ------     i
C DNAREP1_DM#RC        509 AATTTTTCTGGGATATCGATAGATATT--GGGGAATAA------AATGAG 468

  UnnamedSequen        428 AAAATATCATAACATTTTTCAAA-GCGTGGGCGTGACAGCTTTGGGGCGG 476
                               v  ivv  v   ?      - i         i   i      -   
C DNAREP1_DM#RC        467 AAAAAATTTAAAAATTNTTCAAAAGTGTGGGCGTGGCAGTTTTGGG-CGG 419

  UnnamedSequen        477 TTTTTGGGCGTTAGAGTGGGCGTGGCAAA---------------TCGATA 511
                           i   v                        ---------------      
C DNAREP1_DM#RC        418 CTTTAGGGCGTTAGAGTGGGCGTGGCAAAAAGTTTTTTGGCAAATCGATA 369

  UnnamedSequen        512 GAAATTTACAACACCAATACAAAAACGAAAAAATATTAAAACA-TTTTTC 560
                                      v  i    v  ?v i      -   i    v -      
C DNAREP1_DM#RC        368 GAAATTTACAAGACTAATAAAANTATGAAAAA-TATCAAAAAANTTTTTC 320

  UnnamedSequen        561 AAAAGTGTGGGCGTTAGAG---TGGGCG 585
                                         viv  ---      
C DNAREP1_DM#RC        319 AAAAGTGTGGGCGTGGCAGTTTTGGGCG 292

Matrix = 20p43g.matrix
Kimura (with divCpGMod) = 16.51
Transitions / transversions = 0.81 (13/16)
Gap_init rate = 0.07 (14 / 207), avg. gap size = 2.14 (30 / 14)

 2635   7.79 11.86 5.26  UnnamedSequence    533   979    326 C DNAREP1_DM RC/Helitron      1    475    119    
2635 7.79 11.86 5.26 UnnamedSequence 533 979 (326) C DNAREP1_DM#RC/Helitron (119) 475 1 m_b1s001i3

  UnnamedSequen        533 AAAACGAAAAAATATTAAAACATTTTTCAAAAGTGTGGGCGT-------- 574
                               i  i    v   v   v   ?                 --------
C DNAREP1_DM#RC        475 AAAATGAGAAAAAATTTAAAAATTNTTCAAAAGTGTGGGCGTGGCAGTTT 426

  UnnamedSequen        575 ------------------TAGAGTGGGCGTGGCAAAAAGTTGTTTGACAA 606
                           ------------------                       v    i   
C DNAREP1_DM#RC        425 TGGGCGGCTTTAGGGCGTTAGAGTGGGCGTGGCAAAAAGTTTTTTGGCAA 376

  UnnamedSequen        607 GTCGTTAAAAATTTACAACACCAATACAAAAATGAAAAAATATTAAAAAA 656
                           i   v  i          v  i    v  ?v        -   i      
C DNAREP1_DM#RC        375 ATCGATAGAAATTTACAAGACTAATAAAANTATGAAAAA-TATCAAAAAA 327

  UnnamedSequen        657 -TTTTTCAAAAGTGTGGTCGTGGCAATTTTGGGCGGTTTGT-GGC----- 699
                           -                v       i               -   -----
C DNAREP1_DM#RC        326 NTTTTTCAAAAGTGTGGGCGTGGCAGTTTTGGGCGGTTTGTGGGCGTTAG 277

  UnnamedSequen        700 ------CGTGGCAGCATGATTCGACAAACTTGCGCTGCATCTACATATGT 743
                           ------       i    iv  i               i  ----     
C DNAREP1_DM#RC        276 AGTGGGCGTGGCAACATGGGTCAACAAACTTGCGCTGCGTC----TATGT 231

  UnnamedSequen        744 CCCTGGAGTCTGTATGCTTAATCTAAACTTTTTAGCTTTTGTAGTTCCTG 793
                            i  i  i          ?     v   i  i        i         
C DNAREP1_DM#RC        230 CTCTAGAATCTGTATGCTNAATCTCAACCTTCTAGCTTTTATAGTTCCTG 181

  UnnamedSequen        794 AGATCTCAACGTTCATACGGACAGATAGACGGACGGACAGACGGACGGAC 843
                                  i               --------------------       
C DNAREP1_DM#RC        180 AGATCTCGACGTTCATACGGACA--------------------GACGGAC 151

  UnnamedSequen        844 ATGGTTAAATCGACTCGGCTATTGATCCTGATCAAGAATAT--------- 884
                               ii i                                 ---------
C DNAREP1_DM#RC        150 ATGGCCAGATCGACTCGGCTATTGATCCTGATCAAGAATATACGCGCGCG 101

  UnnamedSequen        885 -----ATATACTTTATATGGTCGGAAACGCTTCCTTCTGCCTGTTACATA 929
                           -----                                             
C DNAREP1_DM#RC        100 CGCGNATATACTTTATATGGTCGGAAACGCTTCCTTCTGCCTGTTACATA 51

  UnnamedSequen        930 CTTTTCAACGAATCTAGTATACCCTTTTACTCTACGAGTAACGGGTATAA 979
                                                                             
C DNAREP1_DM#RC         50 CTTTTCAACGAATCTAGTATACCCTTTTACTCTACGAGTAACGGGTATAA 1

Matrix = 20p43g.matrix
Kimura (with divCpGMod) = 7.89
Transitions / transversions = 2.00 (22/11)
Gap_init rate = 0.07 (32 / 446), avg. gap size = 2.44 (78 / 32)
```
