## Supplementary material for "Odorant receptor copy number change, co-expression, and positive selection establish peripheral coding differences between fly species": sup. file 3: RM2sequpload_1628857533.out.html

[ close all hsps ]
[ open all hsps ]

```
                                          position in query-                          -position in repeat-
          %    %    %    query                               C matching   repeat      (left)  end   begin  linkage
+ score  div. del. ins.  sequence         begin  end  (left) + repeat     class/famil begin   end   (left) id/graphic
```

```
+ 1726   13.3 14.3  4.8  UnnamedSequence    115   557  (165) C DNAREP1_DM RC/Helitron    (1)    593      1   1
```

```
ANNOTATION EVIDENCE:
  550  20.26 0.43 18.46  UnnamedSequence    115   344    378 C DNAREP1_DM RC/Helitron    399    593      1    
550 20.26 0.43 18.46 UnnamedSequence 115 344 (378) C DNAREP1_DM#RC/Helitron (1) 593 399 m_b1s001i0

  UnnamedSequen        115 AACAAGAGAGAACGGTGTTGTCGAGTTCCCCGACTATCAGATACCCGTTA 164
                                       i v i v                               
C DNAREP1_DM#RC        593 AACAAGAGAGAATGCTATAGTCGAGTTCCCCGACTATCAGATACCCGTTA 544

  UnnamedSequen        165 CTCGCCCAGTG-GAAGGAGAAGCTGAAATTTCAACCGTTTTTGGATGTTT 213
                              iv i    -   v v   vvi         ---------vi  v   
C DNAREP1_DM#RC        543 CTCAGCTAGTGTGAATGCGAACGCGAAATTTCA---------TAATTTTT 503

  UnnamedSequen        214 TTTGGACGTGGCAAAGAGATTTTTGGCAAATGGAGAGAAGTTTACAAGAC 263
                           i v   ii v --- v    -------- v   i  ------- i  i--
C DNAREP1_DM#RC        502 CTGGGATATCG---ATAGAT--------ATTGGGGA-------ATAAA-- 473

  UnnamedSequen        264 TATAAATCAAAAAATATAAAAACATTATTTAAAAGTGTGGGCGTGGCAGA 313
                           -  i i--       v     --   ?  i                   v
C DNAREP1_DM#RC        472 -ATGAG--AAAAAATTTAAAA--ATTNTTCAAAAGTGTGGGCGTGGCAGT 428

  UnnamedSequen        314 GTTGTGCAGTTTGTGGGCGTTGCAAAATGGG 344
                           v   v  i i  vv       -- i i    
C DNAREP1_DM#RC        427 TTTGGGCGGCTTTAGGGCGTT--AGAGTGGG 399

Matrix = 20p43g.matrix
Kimura (with divCpGMod) = 21.68
Transitions / transversions = 1.00 (19/19)
Gap_init rate = 0.16 (37 / 229), avg. gap size = 1.00 (37 / 37)

 1726  11.19 18.42 0.75  UnnamedSequence    216   557    165 C DNAREP1_DM RC/Helitron      1    402    192    
1726 11.19 18.42 0.75 UnnamedSequence 216 557 (165) C DNAREP1_DM#RC/Helitron (192) 402 1 m_b1s001i1

  UnnamedSequen        216 TGGACGTGGCAAAGAGATTTTTGGCAAATGGAGAGAAGTTTACAAGACTA 265
                              i         i  v            v  v    i            
C DNAREP1_DM#RC        402 TGGGCGTGGCAAAAAGTTTTTTGGCAAATCGATAGAAATTTACAAGACTA 353

  UnnamedSequen        266 -TAAATC---AAAAAATAT-AAAAACATTATTTAAAAGTGTGGGCGTGGC 310
                           -    v?---i        -     v?  v  i                 
C DNAREP1_DM#RC        352 ATAAAANTATGAAAAATATCAAAAAANTTTTTCAAAAGTGTGGGCGTGGC 303

  UnnamedSequen        311 AGAGTTGTGCAGTTTGTGGGCGTT------------GCAAAATGGGTCAA 348
                             vv   v  i             ------------    v         
C DNAREP1_DM#RC        302 AGTTTTGGGCGGTTTGTGGGCGTTAGAGTGGGCGTGGCAACATGGGTCAA 253

  UnnamedSequen        349 CAA------GCTGCGTCT------CTAG---------GTTTAGTC----- 372
                              ------         ------    --------- i ? i  -----
C DNAREP1_DM#RC        252 CAAACTTGCGCTGCGTCTATGTCTCTAGAATCTGTATGCTNAATCTCAAC 203

  UnnamedSequen        373 CTTCTAGCTTTTATAGTTCTTGAGATCTTGACGTTCATACACACGACAGA 422
                                              i        i           i---      
C DNAREP1_DM#RC        202 CTTCTAGCTTTTATAGTTCCTGAGATCTCGACGTTCATACG---GACAGA 156

  UnnamedSequen        423 CGGACAGGACCAAATCGACTCGACTATTGAT----AT------------- 455
                                 v i   i         i        ----  -------------
C DNAREP1_DM#RC        155 CGGACATGGCCAGATCGACTCGGCTATTGATCCTGATCAAGAATATACGC 106

  UnnamedSequen        456 ---CTCTTATATATCCTTAATGTGGTTTGAAACGCTTTCTTCTACCTGTT 502
                           --- v vii?    v   v  i    iv         i     i      
C DNAREP1_DM#RC        105 GCGCGCGCGNATATACTTTATATGGTCGGAAACGCTTCCTTCTGCCTGTT 56

  UnnamedSequen        503 TCATACTTTTCAAAGAATCTAGTATACCCTTTTACTCTACGAGTAACGGG 552
                           v            v                                    
C DNAREP1_DM#RC         55 ACATACTTTTCAACGAATCTAGTATACCCTTTTACTCTACGAGTAACGGG 6

  UnnamedSequen        553 TATAA 557
                                
C DNAREP1_DM#RC          5 TATAA 1

Matrix = 20p43g.matrix
Kimura (with divCpGMod) = 10.71
Transitions / transversions = 1.11 (20/18)
Gap_init rate = 0.04 (14 / 341), avg. gap size = 4.71 (66 / 14)
```
